# Synthetic 5’ UTRs can either up- or down-regulate expression upon RBP binding

**DOI:** 10.1101/174888

**Authors:** Noa Katz, Roni Cohen, Oz Solomon, Beate Kaufmann, Orna Atar, Zohar Yakhini, Sarah Goldberg, Roee Amit

**Affiliations:** Department of Biotechnology and Food Engineering, Technion - Israel Institute of Technology, Haifa 32000, Israel; Department of Computer Science, Technion - Israel Institute of Technology, Haifa 32000, Israel; School of Computer Science, Interdisciplinary Center, Herzeliya 46150, Israel; Russell Berrie Nanotechnology Institute, Technion - Israel Institute of Technology, Haifa 32000, Israel

## Abstract

The construction of complex gene regulatory networks requires both inhibitory and up-regulatory modules. However, the vast majority of RNA-based regulatory “parts” are inhibitory. Using a synthetic biology approach combined with SHAPE-Seq, we explored the regulatory effect of RBP-RNA interactions in bacterial 5’-UTRs. By positioning a library of RNA hairpins upstream of a reporter gene and co-expressing them with the matching RBP, we observed a set of regulatory responses, including translational stimulation, translational repression, and cooperative behavior. Our combined approach revealed three distinct states *in-vivo*: in the absence of RBPs, the RNA molecules can be found either in a molten state that is amenable to translation, or a structured phase that inhibits translation. In the presence of RBPs, the RNA molecules are in a semi-structured phase with partial translational capacity. Our work provides new insight into RBP-based regulation and a blueprint for designing complete gene regulatory circuits at the post-transcriptional level.

## INTRODUCTION

One of the main goals of synthetic biology is the construction of complex gene regulatory networks. The majority of engineered regulatory networks have been based on transcriptional regulation, with only a few examples based on post-transcriptional regulation (Win and Smolke, 2008; Xie et al., 2011; Green et al., 2014; Wroblewska et al., 2015), even-though RNA-based regulatory components have many advantages. Several RNA components have been shown to be functional in multiple organisms (Harvey et al., 2002; Suess et al., 2003; Desai and Gallivan, 2004; Buxbaum et al., 2015; Green et al., 2017). RNA can respond rapidly to stimuli, enabling a faster regulatory response as compared with transcriptional regulation (Hentze et al., 1987; St Johnston, 2005; Saito et al., 2010; Lewis et al., 2017). From a structural perspective, RNA molecules can form a variety of biologically functional secondary and tertiary structures (Green et al., 2014), which enables modularity. For example, distinct sequence domains within a molecule (Khalil and Collins, 2010; Lewis et al., 2017) may target different metabolites or nucleic acid molecules (Werstuck and Green, 1998; Isaacs et al., 2006). All of these characteristics make RNA an appealing target for engineered-based applications (Hutvágner and Zamore, 2002; Rinaudo et al., 2007; Delebecque et al., 2011; Xie et al., 2011; Chen and Silver, 2012; Ausländer et al., 2014; Green et al., 2014; Sachdeva et al., 2014; Pardee et al., 2016).

Perhaps the most well-known class of RNA-based regulatory modules are riboswitches (Werstuck and Green, 1998; Winkler and Breaker, 2005; Henkin, 2008; Wittmann and Suess, 2012; Serganov and Nudler, 2013). Riboswitches are noncoding mRNA segments that regulate the expression of adjacent genes via structural change, effected by a ligand or metabolite. However, response to metabolites cannot be easily used as the basis of a regulatory network, as there is no convenient feedback or feed-forward mechanism for connection with additional network modules. Implementing network modules using RNA-binding proteins (RBPs) could enable an alternative multicomponent connectivity for gene-regulatory networks that is not based solely on transcription factors.

Regulatory networks require both inhibitory and up-regulatory modules. The vast majority of known RBP regulatory mechanisms are inhibitory (Romaniuk et al., 1987; Cerretti et al., 1988; Brown et al., 1997; Schlax et al., 2001; Lim and Peabody, 2002a; Sacerdot Christine et al., 2002). A notable exception is the phage RBP Com, whose binding was demonstrated to destabilize a sequestered ribosome binding site (RBS) of the Mu phage *mom* gene, thereby facilitating translation (Hattman et al., 1991; Wulczyn and Kahmann, 1991). Several studies have attempted to engineer activation modules utilizing RNA-RBP interactions, based on different mechanisms: recruiting the eIF4G1 eukaryotic translation initiation factor to specific RNA targets via fusion of the initiation factor to an RBP (Gregorio et al., 1999; Boutonnet et al., 2004), adopting a riboswitch-like approach (Ausländer et al., 2014), and utilizing an RNA-binding version of the TetR protein (Goldfless et al., 2012). However, despite these notable efforts, RBP-based translational stimulation is still difficult to design in most organisms.

In this study, we employ a synthetic biology reporter assay and *in vivo* SHAPE-Seq (Lucks et al., 2011; Spitale et al., 2013; Flynn et al., 2016) approach to study the regulatory effect controlled by an RBP bound to a hairpin within the 5’ UTR of bacterial mRNA, following a design introduced by (Saito et al., 2010). Our findings indicate that structure-binding RBPs [coat proteins from the bacteriophages GA (Gott et al., 1991), MS2 (Peabody, 1993), PP7 (Lim and Peabody, 2002b), and Qβ (Lim et al., 1996)] can generate a range of translational responses, from previously-observed down-regulation (Saito et al., 2010) to up-regulation. The mechanism for downregulation is most likely steric hindrance of the initiating ribosome by the RBP-mRNA complex. For the 5’ UTR sequences that exhibit up-regulation, RBP binding seems to facilitate a transition from an RNA structure with a low translation rate, into another RNA structure with a higher translation rate. These two experimental features indicate that the up-regulatory elements constitute protein-responsive RNA regulatory elements. Our findings imply that RNA-RBP interactions can provide a platform for constructing gene regulatory networks that are based on translational, rather than transcriptional, regulation.

## RESULTS

### RBP binding can cause either up-regulation or down-regulation

We studied the regulatory effect generated by four RBPs when co-expressed with a reporter construct containing native and non-native binding sites in the 5’ UTR (Fig. 1A). The RBPs used-GCP, MCP, PCP, and QCP-were the coat proteins from the bacteriophages GA, MS2, PP7, and Qβ, respectively (see Table S2). In brief (Fig. 1A and STAR methods), we placed the binding site in the 5’ UTR of the mCherry gene at various positions upstream to the mCherry AUG, induced production of the RBP-mCerulean fusion by addition of N-butyryl-L-homoserine lactone (C_4-_HSL) at 24 different concentrations, and measured both signals (mCherry and mCerulean) to calculate RBP response. An example signal for two duplicates of an up-regulating strain using the mutated PCP binding site PP7-wt positioned at δ=-31 in the 5’ UTR is shown in Fig. 1B. In the upper panel the induction response can be seen for the PCP-mCerulean channel, and in the lower panel the mCherry rate of production for the particular 5’ UTR configuration that results from the induction is shown (see SI for definition).

**Fig. 1.**
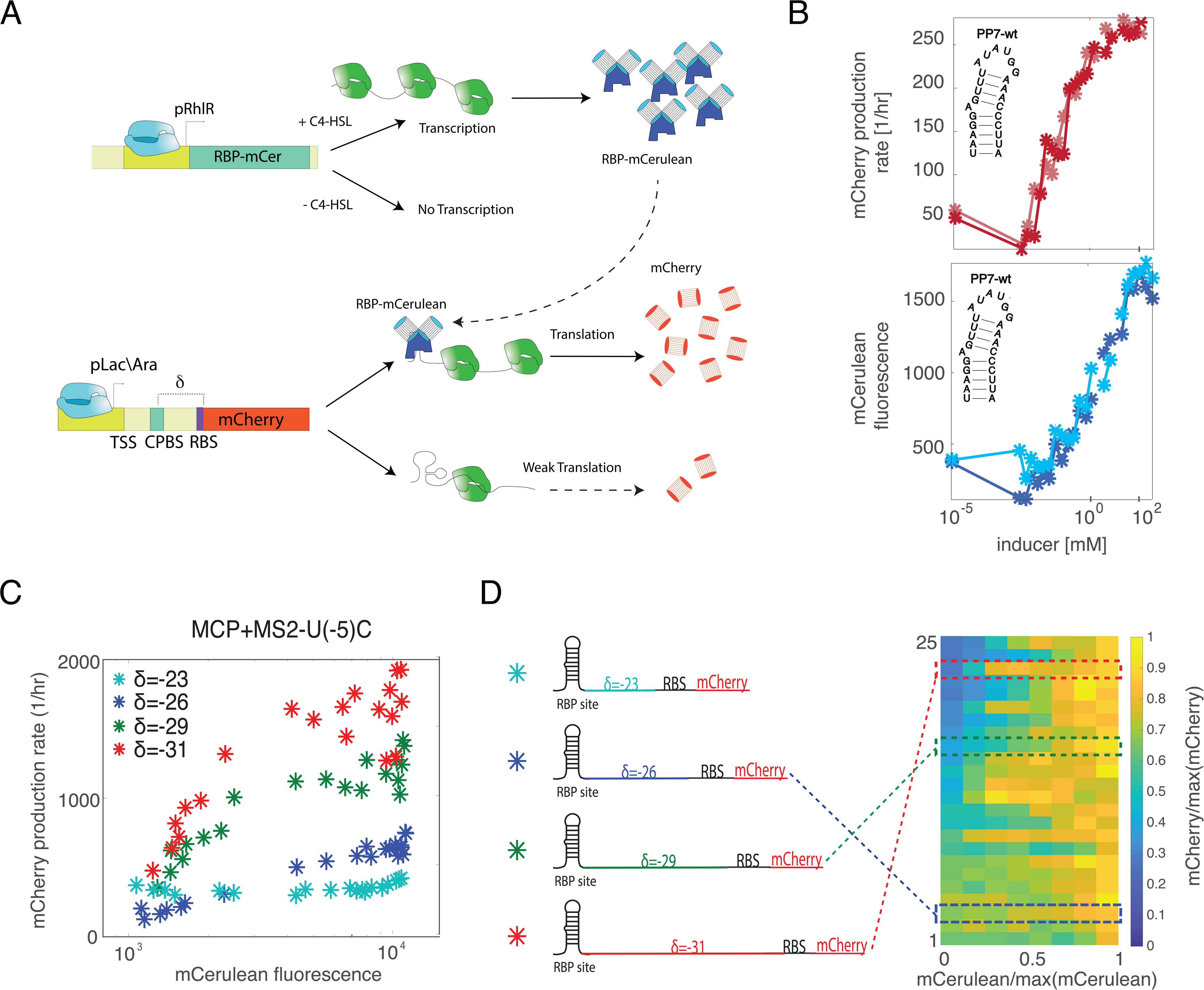
Experimental schematic. (a) Schematic of the experimental system. Top: plasmid expressing the RBP-mCerulean fusion from a pRhlR inducible promoter. Bottom: a second plasmid expressing the reporter mCherry with the RBP binding site encoded within the 5’ end of the gene (at position δ<0). CPBS = coat protein binding site, TSS = transcription start site, RBS = ribosome binding site. (b) A sample data set showing the two fluorescent channels separately for PP7-wt. (Top) mCerulean mean production rate plotted as a function of C_4_-HSL inducer concentration. (Bottom) mCherry reporter expressed from a constitutive pLac/Ara promoter plotted as a function of inducer concentration showing an up-regulatory response which emerges from the RBP-RNA interaction. (c) mCherry production rate for MS2-U(-5)C at four different locations in the 5’ UTR. (d) Illustration of the MS2-U(-5)C site at four different locations (left) and a heatmap of the dose responses for up-regulating variants in the 5’ UTR (right).

To facilitate a more efficient characterization of the dose-response, we analyzed the mCherry production rate for all strains as a function of mCerulean levels. In Fig. 1C (left), we present the sample dose response results for MS2-U(-5)C, together with MCP, at all four different 5’ UTR positions assayed. A sigmoidal response can be observed for three out of the four configurations, with the fold change diminishing as the binding site is positioned closer to the RBS. For the δ=-23 strain, we observe no change in response as a function of the amount of RBP in the cell. To facilitate proper comparison of the regulatory effect across strains, for each strain we opted to normalize both the mCherry rate-of-production and mCerulean expression levels by their respective maximal value for each dose-response function. Such a normalization allows us to properly compare between strains fold-regulation effects, and effective dissociation constant (*K*_*RBP*_), by in effect eliminating the dependence on basal mCherry rate-of-production, and the particular maximal RBP expression levels. Finally, we sorted all normalized dose-responses in accordance with increasing fold up-regulation effect, and plotted the dose-responses obtained in the experiment as a single heatmap facilitating convenient further study and presentation of the data (Fig. 1D).

We constructed our 5’ UTR variants using 11 putative binding sites for the phage coat proteins depicted in Fig. 2A. These structures are based on the three native sites for the RBPs, MS2-wt, PP7-wt, and Qβ-wt (in bold). Different mutations were introduced, some structure-altering, such as the PP7 upper stem short (PP7-USs) and PP7 no-bulge (PP7-nB), and some are structure preserving, such as the MS2-U(-5)C and Qβ-upper stem, lower stem, and loop mutated (Qβ-USLSLm). The minimum free energy of the structure also varies, depending on the kind of mutations introduced. A few mutations in the structure of the binding site can greatly influence the stability of the structure, as is the case for PP7-nB and Qβ-USLSLm.

**Fig. 2.**
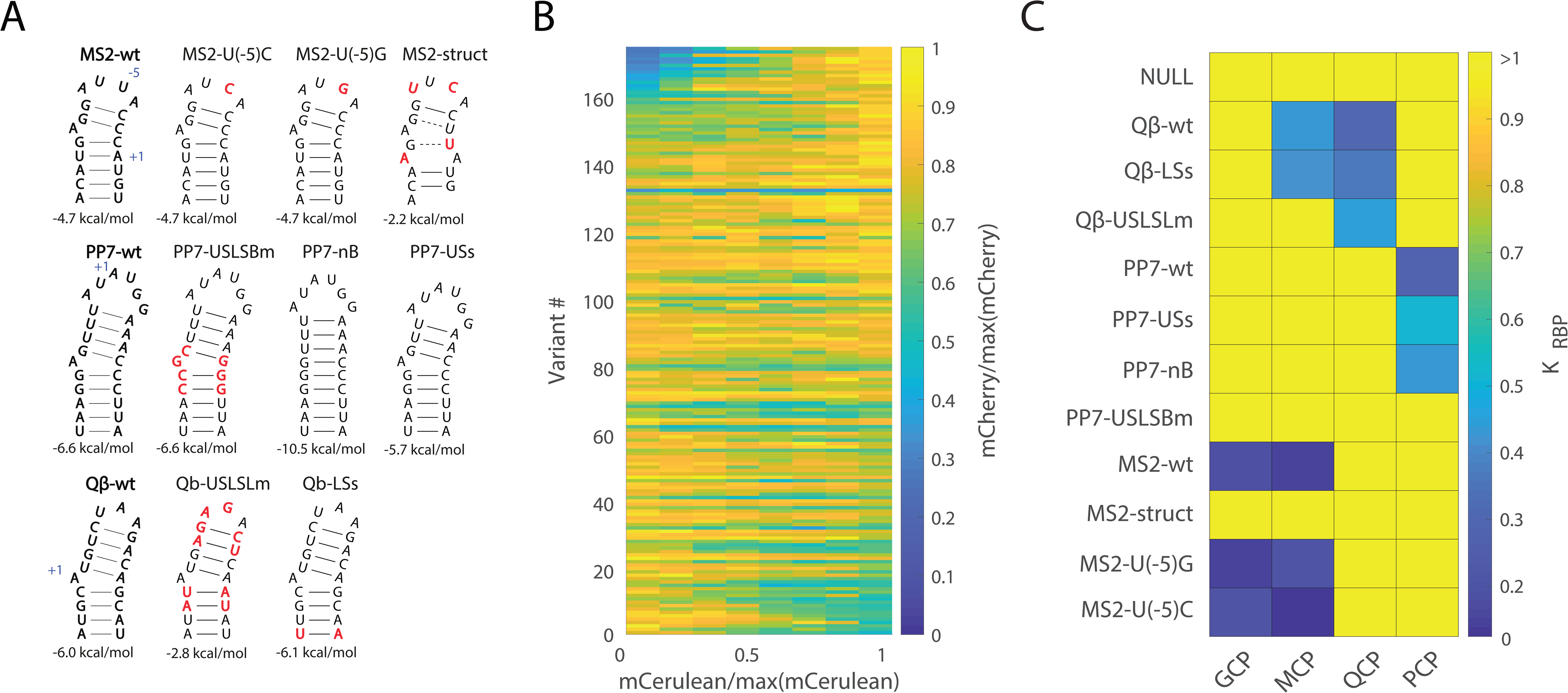
Translational stimulation and repression upon RBP binding in the 5’ UTR. (a) Secondary structure schematic for the 11 binding sites used in the study. Red nucleotides indicate mutations from the original wt binding sequence. Abbreviations: US/LS/L/B = upper stem/ lower stem/ loop/ bulge, m = mutation, s = short, struct = significant change in binding site structure. (b) Heatmap of the dose responses of the 5’ UTR variants. Each response is divided by its maximal mCherry/mCerulean level, for easier comparison. Variants are arranged in order of increasing fold up-regulation. (c) Normalized *K*_*RBP*_ averaged over the different positions. Blue corresponds to low *K*_*RBP*,_ while yellow indicates no binding. If there was no measurable interaction between the RBP and binding site, *K*_*RBP*_ was set to 1. NULL represents no binding site.

We positioned each of the 11 binding sites at three or four different locations upstream of the RBS, that ranged from δ=-21 to δ=-35 nt measured relative to the AUG of the mCherry reporter gene (see Table S1). Altogether, we constructed 44 reporter constructs (including non-hairpin controls), and co-transformed with all four RBPs, resulting in a total of 176 regulatory strains. The normalized and sorted dose-response heatmap for the 5’ UTR constructs for all strains is plotted in Fig. 2B. The dose response functions are arranged in order of increasing fold up-regulation response, with the strongest-repression variants depicted at the bottom. The plot shows that there is a great diversity of responses. We found 24 up-regulating strains (top of the heatmap), 30 down-regulating strains (bottom of the heatmap), and the remaining variants were not found to generate a statistically significant dose-response. A closer examination indicates that the observed repression is generally weak, and at most amounts to about a factor of two reduction from basal levels (turquoise – bottom of the heat map). Notably, the top of the heatmap reveal a moderate up-regulatory dose-response (variant #>140) of up to ∼5-fold, which was not previously observed for these RBPs.

Next, we computed the effective dissociation constant for all dose-responding strains (*K*_*RBP*_), which is defined as the fitted dissociation constant (see STAR methods) normalized by the maximal mCerulean expression level. The resultant *K*_*RBP*_ values obtained for each RBP—binding-site pair are plotted as a heatmap in Fig. 2C. Note, we did not find a position dependence on the values of *K*_*RBP*_ in this experiment (see SI), and thus the values depicted in the heatmap represents an average over multiple 5’ UTR positions. The heatmap shows similar effective dissociation constant values (up to an estimated fit error of 10%) for all binding-site positions, for each of the native binding sites (MS2-wt, PP7-wt, and Qβ-wt), and for the mutated sites with a single mutation (none-structure altering) in the loop region [MS2-U(-5)C and MS2-U(-5)G]. However, for mutated binding sites characterized by small structural deviations from the native structure (PP7-nB and PP7-USs), and for RBPs that bind non-native binding sites (e.g., MCP with Qβ-wt) a higher effective dissociation constant was recorded. Furthermore, deviations in effective dissociation constant were also observed for several of the mutated sites in comparison to a similar measurement that was reported by us recently, when the binding sites were positioned in the ribosomal initiation region (Katz et al., 2018). In particular, both Qβ-USLSLm and Qβ-LSs generated a down-regulatory dose-response signal in the 5’ UTR in the presence of QCP, while no response was detected in the ribosomal initiation region configurations. Conversely, QCP generated a response with the MS2-based sites, MS2-wt and MS2-U(-5)C, in the ribosomal initiation region, while no apparent response was detected when these binding sites were placed in the 5’ UTR. Finally, past *in vitro* studies have recorded a dose-response function for MS2-wt, MS2-U(-5)C, and PP7-USLSBm in the presence of QCP and PCP, while no such effect was observed here for QCP and PCP for any of these sites. Consequently, the nature of the dose response and the mere binding of a protein to a site seems to depend on additional parameters that are not localized solely to the binding site.

### 5’ UTR strains present three translational states

To further study the different types of dose-responses (up-or down-regulation), for each RBP-binding site pair that generate a dose response, we plot the maximal fold-change effect that was recorded over the range of 5’ UTR positions (Fig. 3A). In the panel, we show both maximal down (depicted as fold-values < 1) and up-regulatory dose-response fold changes. The figure shows that the nature of the response does not depend on the RBP, but rather on the binding sites. In particular, both MCP and GCP generate an up-regulatory response for the binding sites MS2-wt, MS2-U(-5)G, and MS2-U(-5)C. Likewise, both MCP and QCP generate a down-regulatory response for Qβ-wt and Qβ-LSs. Conversely, structural mutations that conserve binding (PP7-USs and PP7-nB) can alter the dose-response of PCP from up-regulating (PP7-wt) to down-regulating. Finally, for the case of MS2 with MCP, the size of the fold effect seems to depend on the exact sequence of the binding site. Here, while the native MS2-wt binding site exhibited a maximal fold-change effect of ∼2, a single mutation to the loop region caused the response to increase to a factor of five-fold activation. Taken together, our data indicates that the nature of the response is dependent on the binding site sequence at a single-nucleotide resolution.

**Fig. 3.**
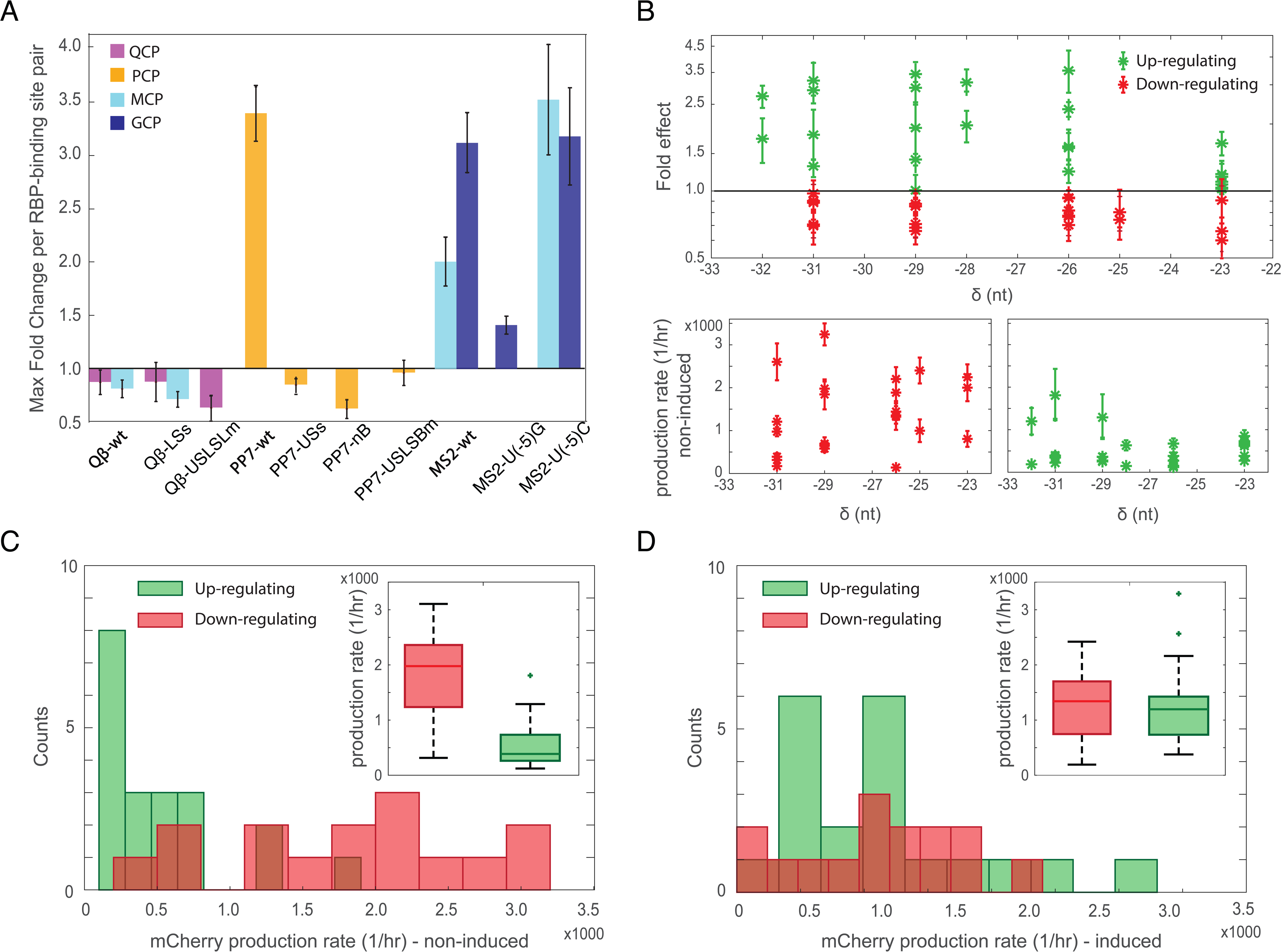
Reporter assay indicates that there may be three distinct translational states (a). Bar graph showing maximal fold change of each RBP—binding-site pair for all 11 binding sites, as follows: QCP-mCerulean (purple), PCP-mCerulean (yellow), MCP-mCerulean (light blue), and GCP-mCerulean (dark blue). Values larger and smaller than one correspond to up-and down-regulation, respectively. The MS2-struct binding site was omitted from the plot due to no observable effect with all RBPs. (b) Top: Fold effect as a function of position for up-regulating strains (green) and down-regulating strains (red). Each point represents a single RBP—binding-site pair. Error bars represent standard deviation from at least two replicates. Bottom: Basal mCherry production rate as a function of position for down-regulating strains (left) and up-regulating strains (right). (c-d). Histograms of mCherry production rate for both regulatory populations, along with matching box-plots (inset) at the non-induced (c) and induced states (d). Mann–Whitney U test (MWW) on the two populations showed a p-value of 1.5702e-04 for the un-induced state, and 0.4822 for the induced state.

We next studied the relationship between the position of the binding site within the 5’ UTR and size of the fold effect. In Fig. 3B top, we plot the fold effect for all RBP-binding site pairs as a function of 5’ UTR position. First, we note that changing the length of the sequence segment downstream to the binding sites does not alter the nature of the dose-response. Second, the plots show that for both the fold repression and fold activation, the effect is mostly unaffected by changing the position of the binding site within the 5’ UTR, except when it is placed in a high proximity to the RBS (position δ = −23), where the activation is diminished. Plots of the basal production rate of both types of strains show a similar picture (Fig. 3B bottom), with the fold activation diminishes as the distance from the RBS is reduced. Next, we compared the absolute rate of production levels between the up-regulating and down-regulating strains, for both the non-induced (Fig. 3C) and fully induced (Fig. 3D) states. For the non-induced states, the mean rate-of-production of the up-regulating strains is around a factor of three less than the mean for the down-regulating strains. Conversely, for the induced state, both distributions converge, and present less than a factor of two difference between the two calculated mean levels. This indicates the translational level associated with the RBP-bound mRNA is similar for all 5’ UTR constructs, independent of the particular binding site or RBP present. Taken together, a picture emerges where there are three main translational states for the 5’ UTR and associated *mCherry* gene, each with its own range of resultant mCherry levels: a closed translationally inactive state occurring for the non-induced up-regulating strains, where the mRNA is predominantly unavailable for translation. An open translationally-active state, which occurs for the non-induced down-regulating strains. Finally, a partially active translational state, which is characterized by an RBP-bound 5’ UTR.

### *In vitro* structural analysis with SHAPE-Seq exhibits a single structural state

Our reporter assay analysis and past results by us and others indicate that there seem to be other factors in play that influence RBP binding and the nature of the dose-response. A prime candidate is the molecular structure that forms *in-vivo* in the presence and absence of the binding protein. This structure is influenced by the sequences which flank the binding site, and the minimum free energy of the hairpin itself. This led us to hypothesize that each state is characterized by a structural fingerprint, which, in turn, is dependent on binding site structure and stability, as well as the flanking sequences. To test our proposed scenario, we chose to focus on two 5’ UTR variants from our library, which encoded the PP7-wt and PP7-USs binding sites, both at δ=-29. In this test case, the entire 5’ UTR is identical for both variants except for a deletion of two nucleotides in the upper stem of the PP7-wt site, which results in the PP7-USs site. This deletion reduces the stability of the PP7-USs binding site (−5.7 kcal/mol) as compared with the native PP7-wt site (−6.6 kcal/mol). First, we wanted to ensure that these variants exhibit the three translational states in their dose response (Fig. 4A). Here, the PP7-wt response function exhibits a low production rate in the absence of induction (∼150 a.u./hr), while rising in a sigmoidal fashion to an intermediate production rate (∼450 a.u./hr) at full induction. For PP7-USs, the basal rate of production level at zero induction is nearly an order of magnitude larger at ∼1100 a.u./hr, and declines gradually upon induction to an intermediate level similar to that observed for PP7-wt.

**Fig. 4.**
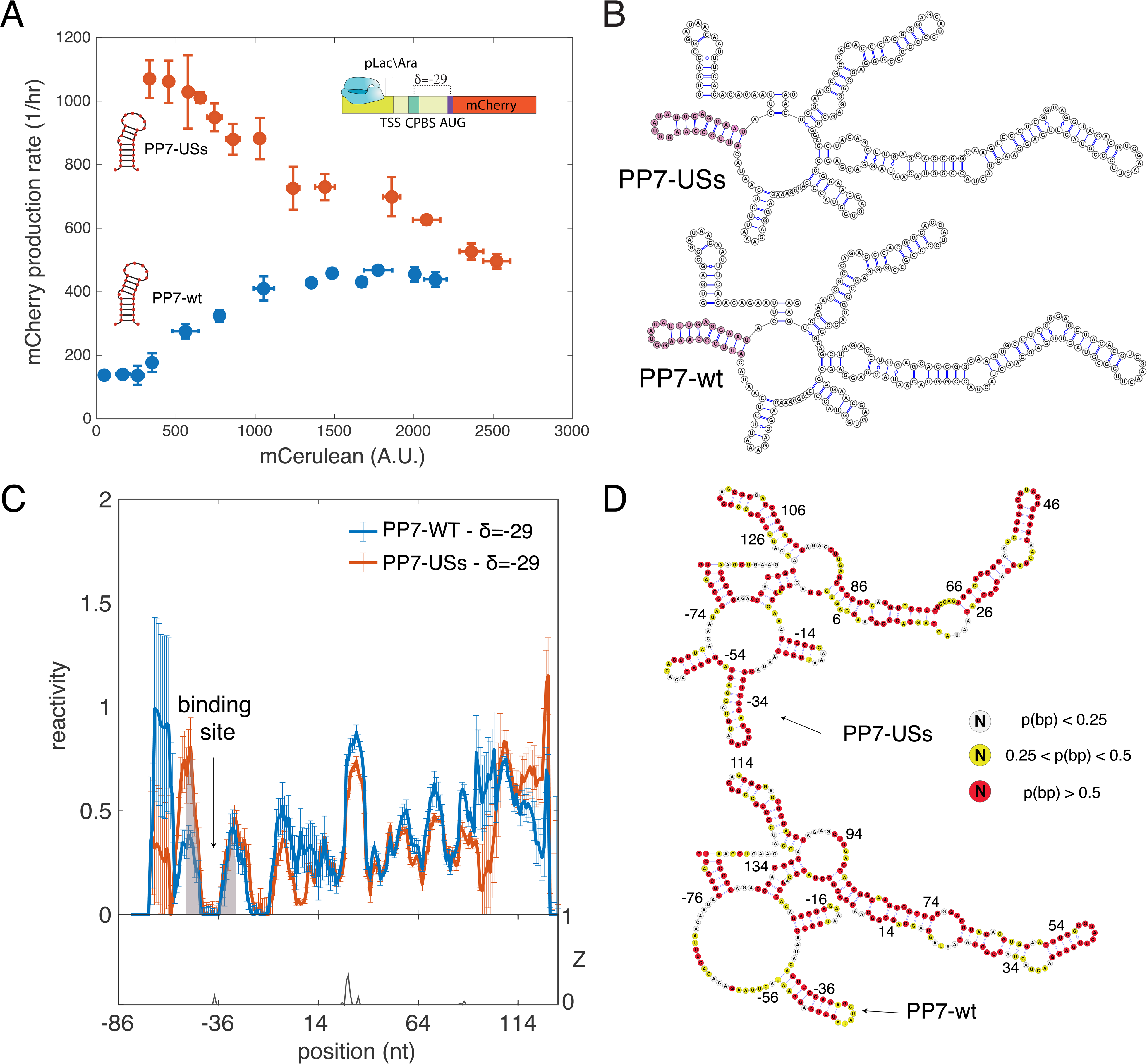
*In vitro* SHAPE-Seq analysis does not reveal two distinct structural states without RBP. (a) Dose response functions for two strains containing the PP7-wt (blue) and PP7-USs (red) binding sites at δ=-29 nt from the AUG. Each data point is an average over multiple mCerulean and mCherry measurements taken at a given inducer concentration. (b) Structure schemes predicted by RNAfold for the 5’ UTR and the first 134 nts of the PP7-wt and PP7-USs constructs (using sequence information only). (c) *In vitro* reactivity analysis for SHAPE-Seq data obtained for two constructs PP7-wt (blue) and PP7-USs (red) at δ=-29. Error-bars are computed using boot-strapping re-sampling of the original modified and non-modified libraries for each strain (see Star Methods) and are also averaged from two biological replicates. The data from the two extra bases for PP7-wt were removed for alignment purposes. (d) Inferred *in vitro* structures for both constructs are constrained by the reactivity scores from (b). Each base is colored by its base pairing probability (red-high, yellow-intermediate, and white-low) calculated based on the structural ensemble via RNAsubopt (Lorenz et al., 2011). Associated with Fig. S1.

Next, we calculated the predicted structure for these two 5’ UTR variants using RNAfold (Hofacker et al., 1994). As expected, the small reduction in binding site stability did not affect the computed structures (Fig. 4B), and both predicted model structures seem identical. Therefore, we chose to directly probe the mRNA structure via SHAPE-Seq. We subjugated the two strains to SHAPE-Seq *in vitro* using 2-methylnicotinic acid imidazole (NAI) suspended in anhydrous dimethyl sulfoxide (DMSO), with DMSO-treated cells as a non-modified control (see Star Methods and Fig. S1 for SHAPE-Seq analysis of 5S-rRNA as positive control). We chose to modify a segment that includes the entire 5’ UTR, and another ∼140 nt of the mCherry reporter gene. In Fig. 4C, we plot the reactivity signals as a function of nucleotide position on the mRNA obtained for both the PP7-wt (blue line) and PP7-USs (red line) constructs at δ=-29 using *in vitro* SHAPE-Seq, after alignment of the two signals (see Star Methods). The reactivity of each base corresponds to the propensity of that base to be modified by NAI (for the definition of reactivity see Star Methods). Both *in vitro* reactivity signals look nearly identical for the entire modified segment of the RNA. This is further confirmed by Z-factor analysis (lower panel), which yields significant distinguishability only for a narrow segment within the coding region (∼+30 nt). We then used the *in vitro* reactivity data to compute the structure of the variants by guiding the computational prediction (Deigan et al., 2009; Ouyang et al., 2013; Washietl et al., 2012; Zarringhalam et al., 2012). In Fig. 4D we show that the SHAPE-derived structures for both constructs are similar to the results of the initial non-constrained RNAfold computation (Fig. 4B), and are nearly indistinguishable from each other. Consequently, the *in vitro* SHAPE-derived structures and reactivity data for the two 5’ UTR variants do not reveal two distinct structural states, which are a precursor for third RBP-bound state.

### *In-vivo* SHAPE-Seq reveals three structural states supporting the three-translation-level hypothesis

Next, we carried out the SHAPE-Seq protocol *in vivo* (see STAR Methods) on induced and non-induced samples for the two variants. In Fig. 5A, we plot the non-induced (RBP-) reactivity obtained for PP7-wt (blue) and PP7-USs (red). The data shows that PP7-USs is more reactive across nearly the entire segment, including all of the 5’ UTR and >50 nt into the coding region. Z-factor analysis reveals that this difference is statistically significant for a large portion of the 5’ UTR and the coding region, suggesting that the PP7-USs is overall more reactive and thus less structured than the PP7-wt fragment. In Fig. 5B, we show that in the induced state (RBP+) both constructs exhibit a weak reactivity signal that is statistically indistinguishable in the 5’ UTR (i.e., Z-factor ∼0 at δ<0). In particular, the region associated with the binding site is unreactive (marked in grey), indicating that the binding site and flanking regions are either protected by the bound RBP, highly structured, or both (see Fig. S2 for further analysis). Consequently, contrary to the *in vitro* SHAPE analysis, for the *in vivo* case the reactivity data for the non-induced case reveals a picture consistent with two distinct translational states, for a sum total of three states when taking the induced reactivity data into account.

**Fig. 5.**
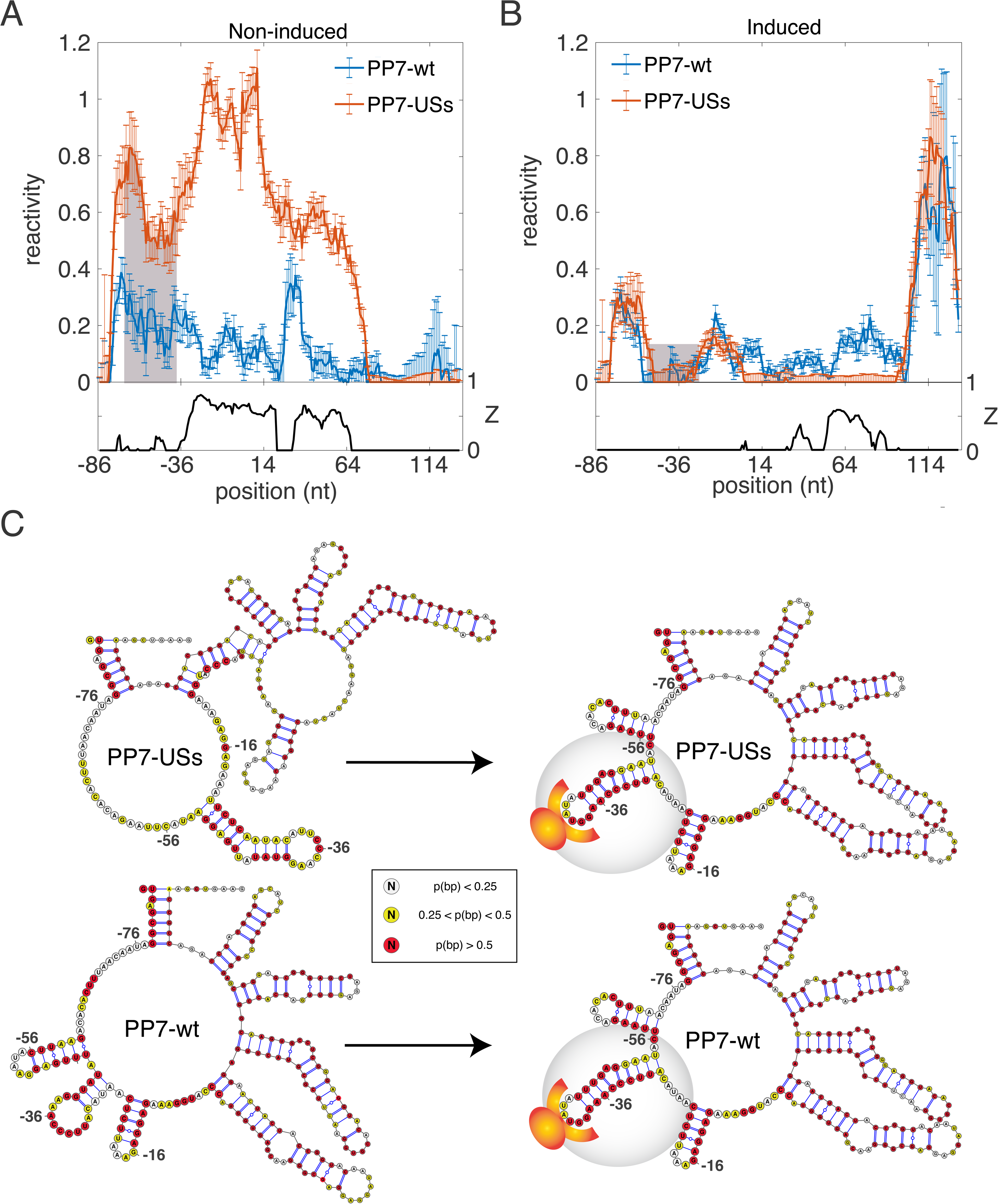
*In vivo* SHAPE-Seq analysis for PP7-wt and PP7-USs strains reveals three structural states. (a-b) Comparison of reactivity analysis computed using *in vivo* SHAPE-Seq data for the non-induced (a) and induced (b) states of PP7-wt (blue) and PP7-USs (red) at δ=-29. Error-bars are computed using boot-strapping re-sampling of the original modified and non-modified libraries for each strain, and also averaged from two biological replicates (see Supp. Information). (c) Inferred *in vivo* structures for all 4 constructs and constrained by the reactivity scores shown in (a-b). Each base is colored by its base pairing probability (red-high, yellow-intermediate, and white-low) calculated based on the structural ensemble via RNAsubopt (Lorenz et al., 2011). For both the PP7-wt and PP7-USs the inferred structures show a distinct structural change in the 5’ UTR as a result of induction of the RBP. Associated with Fig. S2.

To generate additional structural insight, we implemented the constrained structure computation that was used for the *in vitro* samples on the PP7-wt (δ=-29) and PP7-USs (δ=-29) variants (Fig. 5C). In the top schema, we plot the derived PP7-USs non-induced variant, which is non-structured in the 5’ UTR exhibiting a predominantly yellow and white coloring of the individual nucleotide base-pairing probabilities. By contrast, in the PP7-wt non-induced structure (bottom) there are three predicted closely-spaced smaller hairpins that span from −60 to −10 that are predominantly colored by yellow and red except in the predicted loop regions. Both top and bottom structures are markedly different from the *in vitro* structures (Fig. 4D). Neither displays the PP7-wt or PP7-USs binding site, and a secondary hairpin encoding a putative short anti-Shine-Dalgarno (aSD) motif (CUCUU) (Levy et al., 2017), that may partially sequester the RBS, appears only in the PP7-wt non-induced strain. In the induced state, a structure reminiscent of the *in vitro* structure is recovered for both variants with three distinct structural features visible in the 5’ UTR: an upstream flanking hairpin (−72 to −57 for PP7-wt), the binding site (−54 to −30 for PP7-wt), and downstream CUCUU anti-Shine Dalgarno satellite structure (−23 to −10 for both). Taken together, the SHAPE-derived structures for the non-induced and induced strains support three distinct structural configurations for the 5’ UTR, which are consistent with the reporter assay findings and can thus be associated with their respective translational levels.

### Changes to 5’ UTR sequence can alter translational state

We reasoned that we can influence the regulatory response by introducing mutations into the 5’ UTR sequence that can shift the structure from the translationally inactive state to the translationally active state. To do so, we mutated the structure of the flanking sequences in three ways (Fig. 6A): first, by changing the CUCUU motif from the original strains (Fig. 6A-bottom left) into an A-rich segment (Fig. 6A-top right), thus potentially reducing structure formation in the 5’ UTR and potentially shifting the up-regulatory response to a repression effect. Second, by enhancing the aSD motif in the original strains (Fig. 6A-top left), thus encouraging the formation of a structured 5’ UTR and potentially increasing the fold-effect of the up-regulatory strains. Finally, we extended the lower-stem of MS2-wt and PP7-wt binding sites by three, six, and nine base-pairs to increase binding site stability (Fig. 6A-bottom right). We hypothesized that this set of new 5’ UTR variants could help us expand our understanding of the mechanism involved in translational regulation.

**Fig. 6.**
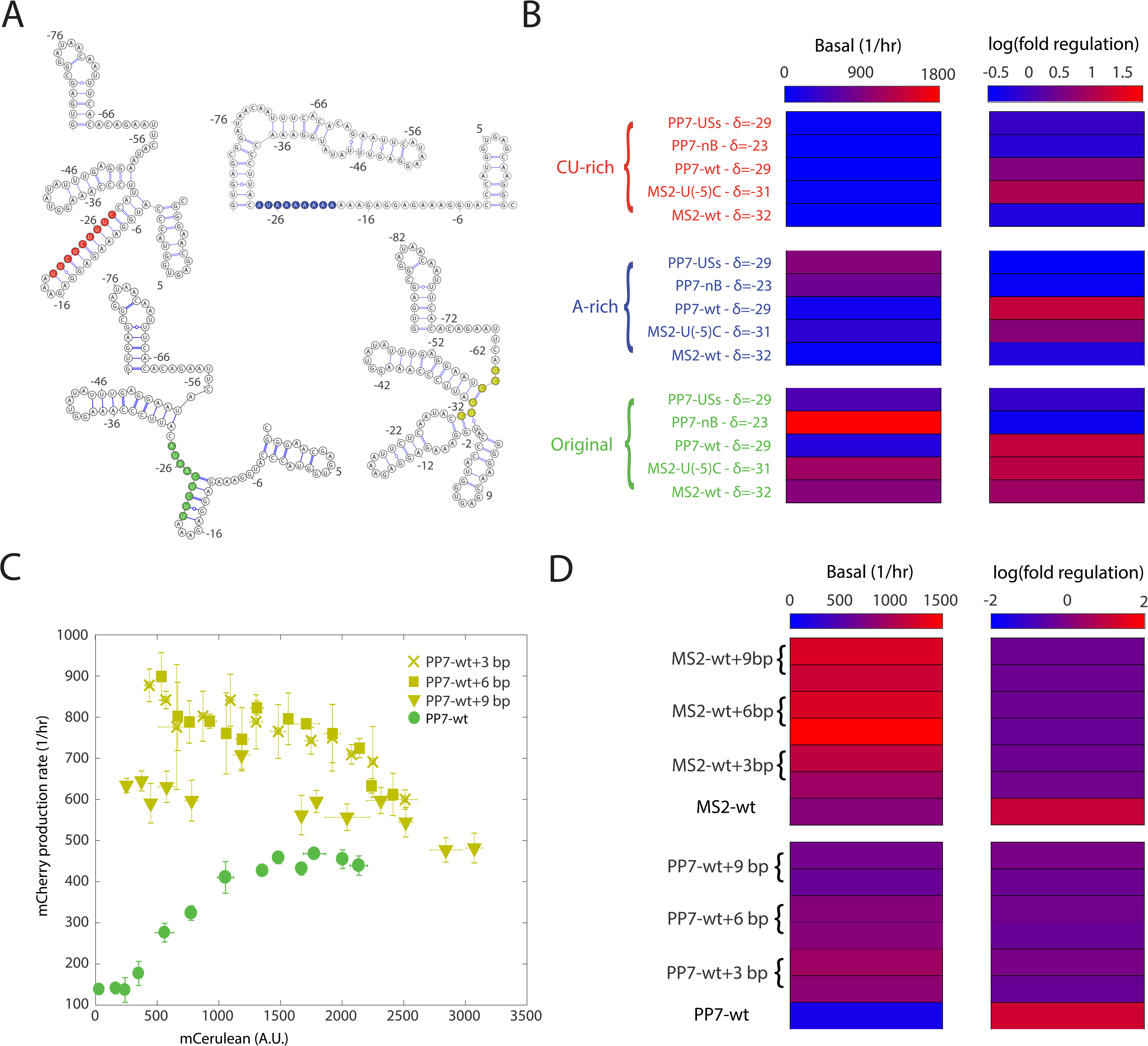
Nature of fold-regulation is dependent on flanking sequences. (a) Schematics for four sample structures computed with RNAfold (using sequence information only), where a short segment of the flanking region to the hairpin was mutated in each strain. Three structures contain the PP7-wt hairpin at δ=-29. (Top-left) CU-rich flanking colored in red. (Top-right) A-rich flanking colored in blue. (Bottom-left) original construct with “random” flanking sequence colored in green. (Bottom-right) PP7-wt hairpin encoded with a longer stem colored in yellow. (b) Variants containing 5 distinct hairpins with either CU-rich (red), A-rich (blue), or original (green) flanking sequences upstream of the RBS. While basal levels are clearly affected by the presence of a strong CU-rich flanking sequence, the nature of the regulatory effect is apparently not determined by the sequence content of the flanking region. (c) Dose response functions for PP7-wt binding sites with an extra 3 (x’s), 6 (squares), and 9 (triangles) stem base-pairs are shown relative to the dose response for PP7-wt (green). (d) Basal levels and logarithm (base 2) of fold change for dose responses of all extended stem constructs with their corresponding RBPs (MCP or PCP). Associated with Fig. S3 and Fig. S4.

First, we synthesized ten additional constructs at δ=-29 with PP7-nB, PP7-USs, PP7-wt, MS2-wt, or MS2-U(-5)C binding sites, in which the sequence between the binding site and the RBS encoded either a strong CU-rich motif, or an A-rich segment (see Table S1). We plot the basal expression level for 15 RBP—binding-site pairs containing the original spacer (green), the spacer with the CU-rich sequence (red), and the A-rich spacer lacking the aSD sequence (blue). The data (Fig.6B – left heatmap) shows that the constructs with a CU-rich flanking region exhibit low basal expression levels as compared with the other constructs, as predicted and previously observed (Levy et al., 2017), while the different A-rich variants do not seem to affect basal expression in a consistent fashion. However, both the up-regulatory and down-regulatory dose responses persist independently of the flanking region content (Fig. 6B-right heatmap – top and middle), as compared with the response recorded for the original flanking sequences (Fig. 6B – right heatmap - bottom).

To check the effect of increasing binding site stability, we designed 6 new variants for the PP7-wt binding sites, by extending the length of the lower stem by three, six, and nine base-pairs, with complementary flanking sequences that are either GU or GC repeats (Fig. S3 and Table 1). When examining the dose-response functions (Fig. 6C-D), the up-regulatory responses was converted to down-regulating responses for all configurations. The basal expression levels for the non-induced state was increased by 3-to 10-fold (Fig. 6D – left heatmap), consistent with the levels previously observed for the non-structured, translationally-active state. Upon induction, the down-regulatory effect that was observed resulted in rate-of-production levels that approached the levels of the original PP7-wt construct at full induction (Fig. 6C), further corroborating the three-state model. Yet, for all stem-extended constructs, the effective dissociation constant increased by 2-to 3-fold (Fig. S3), indicating a potentially weaker binding that may be due to the increased translational activity associated with these constructs. Finally, we checked the effect of temperature on regulation. We studied several strains (RBP-binding sites combinations) in temperatures that ranged from 22°C to 42°C, and found no significant change in regulatory effect for any of the variants studied (Fig. S4). Consequently, it seems that only mutations that are associated with binding site stability seem to affect the state of the non-induced state, whether it will be non-structured and translationally active, or highly structured and translationally inactive.

### A tandem of binding sites can exhibit both cooperativity and complete repression

Finally, to further explore the regulatory potential of the 5’ UTR, we synthesized 28 additional 5’ UTR variants containing two binding sites from our cohort (Fig. 2A), one placed in the 5’ UTR (δ<0), and the other placed in the ribosomal initiation region region (1<δ<15) of the mCherry gene (Fig.7A). In Fig. 7B-D, we plot the dose responses of the tandem variants in the presence of MCP, PCP, and QCP as heatmaps arranged in order of increasing basal mCherry rate of production. Overall, the basal mCherry production rate for all the tandem variants is lower as compared with the single-binding-site variants located in the 5’ UTR. In addition, approximately half of the variants generated a significant regulatory response in the presence of the RBP, while the other half seem to be repressed at the basal level, with no RBP-related effect detected.

**Fig. 7.**
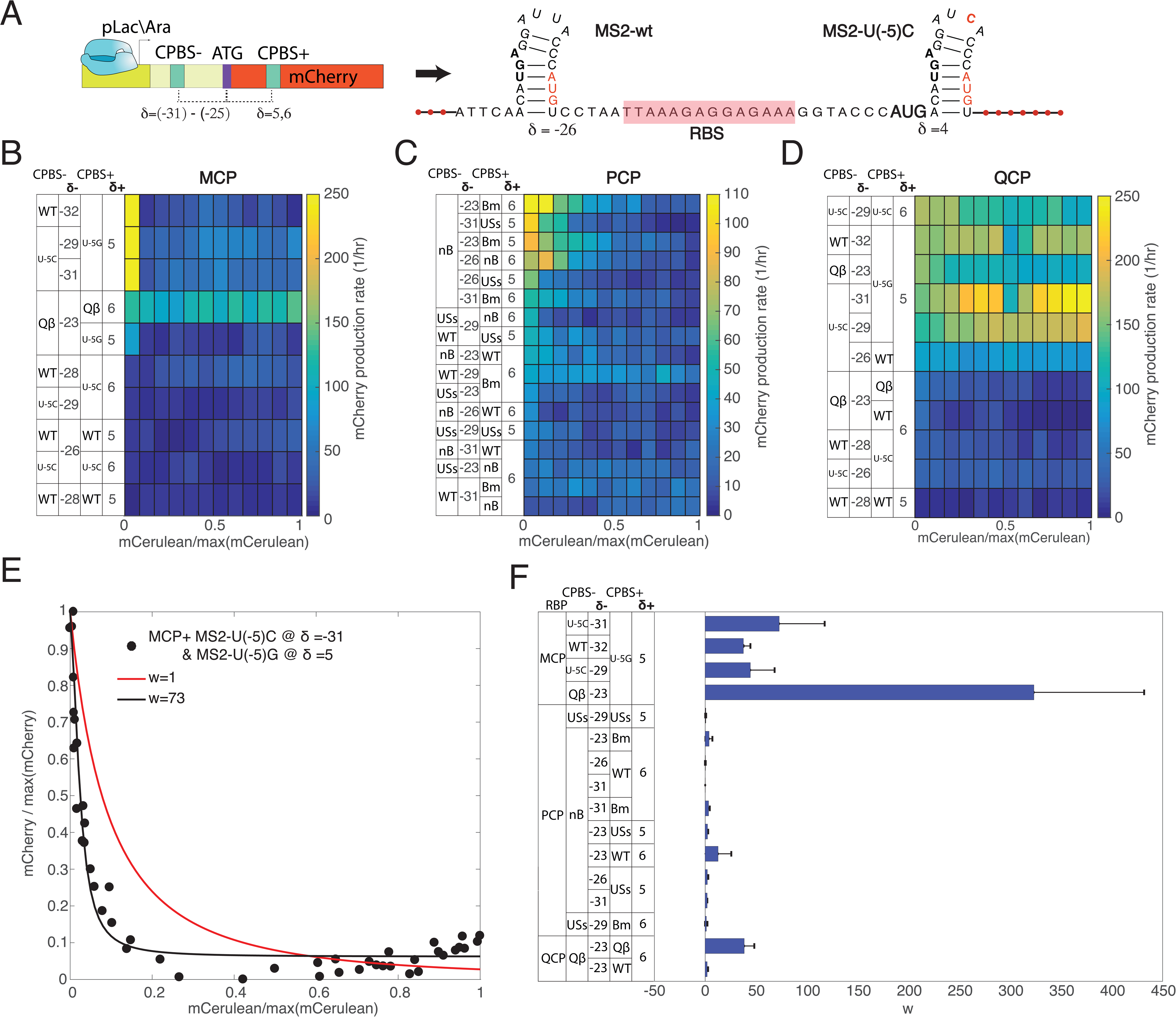
mRNAs with a tandem of hairpins. (a) Schematic of the mRNA molecules with a single binding site at the 5’ UTR (δ<0) and a single binding site in the gene-header region (δ>0). Extra bases were added downstream to the binding site where necessary to retain the open reading frame. (b-d) Heatmap corresponding the dose-response function observed for MCP (b), PCP (c) and QCP (d). In all heatmaps, the dose-response is arranged in order of increasing mCherry rate of production, with the lowest-expressing variant at the bottom. The binding-site abbreviations are as follows: For MCP (b) and QCP (d), WT is MS2-wt, U(-5)G is MS2-U(-5)G, U(-5)C is MS2-U(-5)C, and Qβ is Qβ-wt. For PCP (c), WT is PP7-wt, nB is PP7-nB, Bm is PP7-LSLSBm, and USs is PP7-USs. (e) A sample fit using the cooperativity model (see Supp. Information). (f) Bar plot depicting the extracted cooperativity factors *w* for all the tandems that displayed either an up-or down-regulatory effect. Associated with Fig. S5 and Fig. S6.

For MCP (Fig. 7B), we observed strong repression for four of the ten variants tested, with the MS2-U(-5)G binding site positioned in the ribosomal initiation region for all four repressed variants. With different ribosomal initiation region binding sites [MS2-wt, Qβ-wt, or MS2-U(-5)C], basal mCherry rate of production was reduced to nearly zero. For PCP (Fig. 7C), a similar picture emerges, with several variants exhibiting a strong dose-response repression signature, while no regulatory effect was observed for others. In terms of basal mCherry production rate, the variants in the top six all encode the PP7-nB binding site in the 5’ UTR. Moreover, all eight variants with a PP7-nB positioned in the 5’ UTR exhibit a down-regulatory response. These observations are consistent with the data shown in Fig. 6C-D, where the binding sites with longer stems resulted in larger basal mCherry rate of production, presumably due to increased hairpin stability. For other PP7 binding-site combinations a lower basal level, and hence lower fold-repression effect, is observed.

In Fig. 7D, we present the dose-response heatmaps obtained for QCP. Here, we used the same tandem variants as for MCP, due to the binding cross-talk between both proteins shown in (Fig. 1B). Notably, the dose responses for these tandems in the presence of QCP vary substantially as compared with that observed for MCP. While the site MS2-U(-5G) is still associated with higher basal expression when positioned in the ribosomal initiation region, only three variants (as compared with five for MCP) do not seem to respond to QCP. In particular, two variants, each containing MS2-U(-5)G in the ribosomal initiation region and MS2-U(-5)C in the 5’ UTR, exhibit a 2-fold up-regulatory dose response, as compared with a strong down-regulatory effect for MCP. Given the propensity of binding sites in the ribosomal initiation region to generate a strong repression effect (Katz et al., 2018), the up-regulatory effect observed here is consistent with a lack of binding of QCP to MS2-U(-5)G in the ribosomal initiation region (as was observed before), thus facilitating the up-regulation effect that was observed previously for MCP with MS2-U(-5)C in the single binding site strains.

Finally, we measured the effective cooperativity factor *w* (see Fig.S5 for fitting model) for repressive tandem constructs in the presence of their corresponding cognate RBPs. In Fig. 7E, we plot a sample fit for a MS2-U(-5)C/MS2-U(-5)G tandem in the presence of MCP. The data shows that when taking into account the known *K*_*RBP*_ values that were extracted for the single binding site variants, a fit with no cooperativity (*w=*1) does not explain the data well (red line). However, when the cooperativity parameter is not fixed, a good description for the data is obtained for 50*<w<*80 (best fit at *w*=73). In Fig. 7F we plot the extracted cooperativity parameter for each of the 16 tandems displaying a regulatory response with calculated *K*_*RBP*_ values for both sites (see Fig. S5 and Table S5 for fits and parameter values, respectively). Altogether, at least 6 of the 16 tandems exhibited strong cooperative behavior. For MCP and QCP five of the six relevant tandems displayed strong cooperativity (*w*>25). For PCP, only two of the ten tandems displayed weak cooperativity (1<*w*<25). These tandems had less than 30 nt between the two PCP binding sites.

The cooperative behavior, which reflects overall increase in affinity of the RBP to the molecule when there is more than one binding site present, may also indicate increased stability of the hairpin structures. An increased stability can explain two additional features of the tandems, that were not observed for the single binding site constructs: the QCP up-regulatory response observed for the MS2-U(-5)C/MS2-U(-5)G tandem, and the decreased basal mCherry rate of production levels. Overall, the effective dissociation constant of the tandem-and single-binding-site constructs together with the RBPs can be varied over a range of specificities that spans approximately an order of magnitude, depending on the chosen 5’ UTR and gene-header sequences.

## DISCUSSION

In recent years, synthetic biology approaches have been increasingly used to map potential regulatory mechanisms of transcriptional and translational regulation, in both eukaryotic and bacterial cells (Kinney et al., 2010; Sharon et al., 2012; Dvir et al., 2013; Weingarten-Gabbay et al., 2016; Peterman and Levine, 2016; Levy et al., 2017). Here, we built on the design introduced by (Saito et al., 2010) to explore the regulatory potential of RBP-RNA interactions in bacterial 5’ UTRs, using a synthetic biology approach combined with the SHAPE-Seq method. Using a library of RNA variants, we found a complex set of regulatory responses, including translational repression, translational stimulation, and cooperative behavior. The up-regulation phenomenon, or translational stimulation, had been reported only once for a single natural example in bacteria, yet was mimicked here by all four RBPs at multiple 5’ UTR positions.

Our expression level data on the single-binding-site constructs hints that the mechanism which drives the complexity observed can be described by a three-state system. Using both the SHAPE-Seq experiment and the reporter assay, we found a translationally-active and weakly-structured 5’ UTR state, a translationally-inactive and highly-structured 5’ UTR state, and an RBP-bound state with partial translation capacity. As a result, the same RBP can either up-regulate or down-regulate expression, depending on 5’ UTR sequence context. This description deviates from the classic two-state regulatory model, which is often used as a theoretical basis for describing transcriptional and post-transcriptional regulation (Bintu et al., 2005). In a two-state model, a substrate can either be bound or not bound by a ligand, leading to either an active or inactive regulatory state. This implies that in the two-state scenario, a bound protein cannot both be an “activator” and a “repressor” without an additional interaction or constraint which alters the system.

The appearance of two distinct mRNA states in the non-induced case *in vivo*, as compared with only one *in vitro*, suggests that *in vivo* the mRNA molecules can fold into one of two distinct phases: a molten phase that is amenable to translation, and a structured phase that inhibits translation. A previous theoretical study by Schwab and Bruinsma (SB) (Schwab and Bruinsma, 2009) showed that a first-order phase transition separating a molten and a structured phase for mRNA can occur, if a strong attractive interaction between the non-base-paired segments of the molecule exists within the system (see Fig. S6). Such an interaction destabilizes the base-pairing of branched structures, and if sufficiently strong leads to complete melting of the molecule into a non-structured form. It is possible that such attractive interaction between non-base-paired segments is mediated by the ribosome, which is known to destabilize base-paired structures during translation.

Furthermore, the RBP-bound states which yielded indistinguishable *in vivo* SHAPE-Seq data together with a convergence of the induced up-and down-regulating expression distributions, is also consistent with the SB model. In this case, the SB phase diagram (see Fig. S6) shows that a weaker attractive interaction does not yield a first order phase transition, but rather a continuous transition from a fully-structured phase through a partially-structured phase to the fully-molten state. Since the bound RBP stabilizes the hairpin structure, counteracting the destabilizing effect of the ribosome, in the context of the SB model this effect may lead to a reduction in the strength of the “attractive” interaction. Therefore, it is possible that this binding event shifts the RNA molecules into the portion of the phase diagram (see Fig. S6 - bottom) in which the partially folded state minimizes the free energy, leading to the observed expression level and reactivity measurements in the induced phase.

Our work presents an important step in understanding and engineering post-transcriptional regulatory networks. Throughout this paper we attempted at increasing the synthetic biology utility of our work, the highlight being the direct activation of translation via a single RBP—binding-site pair. As a result, our synthetic regulatory modules can be viewed as a new class of “protein-sensing-riboswitches”, which, given the hypothesized phase-based characterization, may ultimately have a wide utility in gene regulatory applications. Together with our previous work of positioning the sites in the ribosomal initiation region (Katz et al., 2018), we offer a set of modestly up-regulating and a range of down-regulating RBP-binding site pairs with tuneable affinities for four RBPs, three of which are orthogonal to each other (PCP, GCP, and QCP). While we emphasize that our results were obtained in *E. coli*, given the propensity of RBPs to alter the RNA structure via direct interaction, it is tempting to speculate that such an interaction may be a generic 5’ UTR mechanism that could be extended to other RBPs and other organisms.

How difficult is it to design an up-regulatory dose-response for an RBP *de novo*? Unfortunately, our data does not provide a satisfactory mechanistic outcome for a quantitative prediction, but a qualitative phase-based description, which is an initial step. Our experiments revealed no particular structural features that were associated with this regulatory switch, such as the release of a sequestered RBS, which has been reported before as a natural mechanism for translational stimulation (Hattman et al., 1991; Wulczyn and Kahmann, 1991). Moreover, attempting to allocate a structural state for a certain sequence *in vivo* using *in-silico* RNA structure prediction tools is not a reliable approach, due to mechanistic differences between the *in vivo* and *in vitro* environment, which these models understandably do not take into account. Therefore, to provide a predictive blueprint for which sequences are likely to be translationally inactive in their native RBP unbound state, a better understanding of both RNA dynamics and the interaction of RNA with the translational machinery *in-vivo* needs to be established. Yet, our findings suggest that generating translational stimulation using RBPs may not be as difficult as previously thought. At present, the best approach to designing functional elements is to first characterize experimentally a small library of a variety of designs, and subsequently select and optimize the variants that exhibit interesting functionality. Finally, the described constructs add to the growing toolkit of translational regulatory parts, and provide a working design for further exploration of both natural and synthetic post-transcriptional gene regulatory networks.

## SUPPLEMENTARY DATA

6 Supplementary Figures.

5 Supplementary Tables.

Supplementary Data are available at Cell Systems online.

## ACKNOWLEDGEMENT

The authors would like to acknowledge the Technion’s LS&E staff (Tal Katz-Ezov and Anastasia Diviatis) for help with sequencing.

## AUTHOR CONTRIBUTIONS

NK designed and carried out the expression level experiments and analysis for all constructs. NK also conducted several of the SHAPE-seq experiments with RC. RC and BK designed and carried out the SHAPE-Seq experiments. OS and ZY helped analyse the SHAPE-Seq data. SG and OA assisted and guided the experiments and analysis. RA supervised the study. NK, RA, SG, and BK wrote the manuscript.

## STAR * METHODS

### CONTACT FOR REAGENT AND RESOURCE SHARING

Further information and requests for reagents may be directed to and will be fulfilled by the corresponding author Roee Amit.

### EXPERIMENTAL MODEL AND SUBJECT DETAILS

*E. coli* TOP10 cells were obtained from Invitrogen, cat number C404006 (see also Key resource table). Cells were grown in Laural Broth (LB) with appropriate antibiotics overnight at 37 °C and 250 rpm. In the morning, they were diluted by a factor of 100 to semi-poor medium (SPM) consisting of 95% bio-assay (BA) and 5% LB with appropriate antibiotics and different inducer concentrations at 37 °C and 250 rpm for 1hr to 4hrs (Methods Details section for more details).

## METHOD DETAILS

### Design and construction of binding-site plasmids

Binding-site cassettes (see Table S1) were ordered as double-stranded DNA minigenes from either Gen9 or Twist Bioscience. Each minigene was ∼500□bp long and contained the following parts: Eagl restriction site, ∼40 bases of the 5’ end of the Kanamycin (Kan) resistance gene, pLac-Ara constitutive promoter, ribosome binding site (RBS), and a KpnI restriction site. In addition, each cassette contained one or two wild-type or mutated RBP binding sites, either upstream or downstream to the RBS (see Table S1), at varying distances. All binding sites were derived from the wild-type binding sites of the coat proteins of one of the four bacteriophages GA, MS2, PP7, and Qβ. For insertion into the binding-site plasmid backbone, minigene cassettes were double-digested with Eagl-HF and either KpnI or ApaLI (New England Biolabs [NEB]). The digested minigenes were then cloned into the binding-site backbone containing the rest of the mCherry gene, terminator, and the remainder of the Kanamycin resistance gene, by ligation and transformation into *E. coli* TOP10 cells (ThermoFisher Scientific). All the plasmids were sequence-verified by Sanger sequencing. Purified plasmids were stored in 96-well format, for transformation into *E. coli* TOP10 cells containing one of the four fusion-RBP plasmids (see below).

### Design and construction of fusion-RBP plasmids

RBP sequences lacking a stop codon were amplified via PCR off either Addgene or custom-ordered templates (Genescript or IDT, see Table S2). All RBPs presented (GCP, MCP, PCP, and QCP) were cloned into the RBP plasmid between restriction sites KpnI and AgeI, immediately upstream of an mCerulean gene lacking a start codon, under the so-called RhlR promoter [containing the *rhlAB* las box (Medina et al., 2003)] and induced by N-butyryl-L-homoserine lactone (C_4_-HSL). The backbone contained an Ampicillin (Amp) resistance gene. The resulting fusion-RBP plasmids were transformed into *E. coli* TOP10 cells. After Sanger sequencing, positive transformants were made chemically-competent and stored at −80°C in 96-well format.

### Transformation of binding-site plasmids

Binding-site plasmids stored in 96-well format were simultaneously transformed into chemically-competent bacterial cells containing one of the fusion plasmids, also prepared in 96-well format. After transformation, cells were plated using an 8-channel pipettor on 8-lane plates containing LB-agar with relevant antibiotics (Kan and Amp). Double transformants were selected, grown overnight, and stored as glycerol stocks at −80°C in 96-well plates.

### Single clone expression level assay

Dose-response fluorescence experiments were performed using a liquid-handling system in combination with a Liconic incubator and a TECAN Infinite F200 PRO platereader. Each measurement was carried out in duplicates. Double-transformant strains were grown at 37°C and 250 rpm shaking in 1.5 ml LB in 48-well plates with appropriate antibiotics (Kan and Amp) over a period of 16 hours (overnight). In the morning, the inducer for the rhlR promoter C_4_-HSL was pipetted manually to 4 wells in an inducer plate, and then diluted by the robot into 24 concentrations ranging from 0 to 218 nM. While the inducer dilutions were being prepared, semi-poor medium consisting of 95% bioassay buffer (for 1 L: 0.5 g Tryptone [Bacto], 0.3 ml Glycerol, 5.8 g NaCl, 50 ml 1M MgSO4, 1ml 10xPBS buffer pH 7.4, 950 ml DDW) and 5% LB was heated in the incubator, in 96-well plates. The overnight strains were then diluted by the liquid-handling robot by a factor of 100 into 200 μL of pre-heated semi-poor medium, in 96-well plates suitable for fluorescent measurement. The diluted inducer was then transferred by the robot from the inducer plate to the 96-well plates containing the strains. The plates were shaken at 37°C for 6 hours. Note, that induction was only used for the rhlR promoter, which controls the expression of the RBP-mCerulean fusion. The pLac/Ara promoter controlling the mCherry reporter gene functioned as a constitutive promoter of suiTable Strength in our strains and did not require IPTG or Arabinose induction.

Measurement of OD, and mCherry and mCerulean fluorescence were taken via a platereader every 30 minutes. Blank measurements (growth medium only) were subtracted from all fluorescence measurements. For each day of experiment (16 different strains), a time interval of logarithmic growth was chosen (T_0_ to T_final_) according to the measured growth curves, between the linear growth phase and the stationary (T_0_ is typically the third measured time point). Six to eight time points were taken into account, discarding the first and last measurements to avoid errors derived from inaccuracy of exponential growth detection. Strains that showed abnormal growth curves or strains where logarithmic growth phase could not be detected, were not taken into account and the experiment was repeated. See Fig. S2 for experimental schematic and a sample data set.

### SHAPE-Seq experimental setup

LB medium supplemented with appropriate concentrations of Amp and Kan was inoculated with glycerol stocks of bacterial strains harboring both the binding-site plasmid and the RBP-fusion plasmid and grown at 37°C for 16 hours while shaking at 250 rpm. Overnight cultures were diluted 1:100 into SPM. Each bacterial sample was divided into a non-induced sample and an induced sample in which RBP protein expression was induced with 250 nM N-butanoyl-L-homoserine lactone (C_4_-HSL), as described above.

Bacterial cells were grown until OD_600_=0.3, 2 ml of cells were centrifuged and gently resuspended in 0.5 ml SPM. For *in vivo* SHAPE modification, cells were supplemented with a final concentration of 30 mM 2-methylnicotinic acid imidazole (NAI) suspended in anhydrous dimethyl sulfoxide (DMSO, Sigma Aldrich) (Spitale et al., 2013), or 5% (v/v) DMSO. Cells were incubated for 5 min at 37°C while shaking and subsequently centrifuged at 6000 g for 5 min. RNA isolation of 5S rRNA was performed using TRIzol-based standard protocols. Briefly, cells were lysed using Max Bacterial Enhancement Reagent followed by TRIzol treatment (both from Life Technologies). Phase separation was performed using chloroform. RNA was precipitated from the aqueous phase using isopropanol and ethanol washes, and then resuspended in RNase-free water. For the strains harboring PP7-wt δ=-29 and PP7-USs δ=-29, column-based RNA isolation (RNeasy mini kit, QIAGEN) was performed. Samples were divided into the following sub-samples (except for 5S rRNA, where no induction was used):

1. induced/modified (+C_4_-HSL/+NAI)
2. non-induced/modified (-C_4_-HSL/+NAI)
3. induced/non-modified (+C_4_-HSL/+DMSO)
4. non-induced/non-modified (-C_4_-HSL/+DMSO).

*In vitro* modification was carried out on DMSO-treated samples (3 and 4) and has been described elsewhere (Flynn et al., 2016). 1500 ng of RNA isolated from cells treated with DMSO were denatured at 95°C for 5 min, transferred to ice for 1 min and incubated in SHAPE-Seq reaction buffer (100 mM HEPES [pH 7.5], 20 mM MgCl_2_, 6.6 mM NaCl) supplemented with 40 U of RiboLock RNAse inhibitor (Thermo Fisher Scientific) for 5 min at 37°C. Subsequently, final concentrations of 100 mM NAI or 5% (v/v) DMSO were added to the RNA-SHAPE buffer reaction mix and incubated for an additional 5 min at 37°C while shaking. Samples were then transferred to ice to stop the SHAPE-reaction and precipitated by addition of 3 volumes of ice-cold 100% ethanol, followed by incubation at −80°C for 15 min and centrifugation at 4°C, 17000 g for 15 min. Samples were air-dried for 5 min at room temperature and resuspended in 10 µl of RNAse-free water.

Subsequent steps of the SHAPE-Seq protocol, that were applied to all samples, have been described elsewhere (Watters et al., 2016), including reverse transcription (steps 40-51), adapter ligation and purification (steps 52-57) as well as dsDNA sequencing library preparation (steps 68-76). 1000 ng of RNA were converted to cDNA using the reverse transcription primers (for details of primer and adapter sequences used in this work see Table S3) for mCherry (#1) or 5S rRNA (#2) that are specific for either the mCherry transcripts (PP7-USs δ=-29, PP7-wt δ=-29). The RNA was mixed with 0.5 µM primer (#1) or (#2) and incubated at 95°C for 2 min followed by an incubation at 65°C for 5 min. The Superscript III reaction mix (Thermo Fisher Scientific; 1x SSIII First Strand Buffer, 5 mM DTT, 0.5 mM dNTPs, 200 U Superscript III reverse transcriptase) was added to the cDNA/primer mix, cooled down to 45°C and subsequently incubated at 52°C for 25 min. Following inactivation of the reverse transcriptase for 5 min at 65°C, the RNA was hydrolyzed (0.5 M NaOH, 95°C, 5 min) and neutralized (0.2 M HCl). cDNA was precipitated with 3 volumes of ice-cold 100% ethanol, incubated at −80°C for 15 minutes, centrifuged at 4°C for 15 min at 17000 g and resuspended in 22.5 µl ultra-pure water. Next, 1.7 µM of 5’ phosphorylated ssDNA adapter (#3) (see Table S3) was ligated to the cDNA using a CircLigase reaction mix (1xCircLigase reaction buffer, 2.5 mM MnCl_2_, 50 µM ATP, 100 U CircLigase). Samples were incubated at 60°C for 120 min, followed by an inactivation step at 80°C for 10 min. cDNA was ethanol precipitated (3 volumes ice-cold 100% ethanol, 75 mM sodium acetate [pH 5.5], 0.05 mg/mL glycogen [Invitrogen]). After an overnight incubation at −80°C, the cDNA was centrifuged (4°C, 30 min at 17000 g) and resuspended in 20 µl ultra-pure water. To remove non-ligated adapter (#3), resuspended cDNA was further purified using the Agencourt AMPure XP beads (Beackman Coulter) by mixing 1.8x of AMPure bead slurry with the cDNA and incubation at room temperature for 5 min. The subsequent steps were carried out with a DynaMag-96 Side Magnet (Thermo Fisher Scientific) according to the manufacturer’s protocol. Following the washing steps with 70% ethanol, cDNA was resuspended in 20 μl ultra-pure water and were subjected to PCR amplification to construct dsDNA library as detailed below.

### SHAPE-Seq library preparation and sequencing

To produce the dsDNA for sequencing 10ul of purified cDNA from the SHAPE procedure (see above) were PCR amplified using 3 primers: 4nM mCherry selection (#4) or 5S rRNA selection primer (#5), 0.5µM TruSeq Universal Adapter (#6) and 0.5µM TrueSeq Illumina indexes (one of #7-26) (Table S3) with PCR reaction mix (1x Q5 HotStart reaction buffer, 0.1 mM dNTPs, 1 U Q5 HotStart Polymerase [NEB]). A 15-cycle PCR program was used: initial denaturation at 98°C for 30 s followed by a denaturation step at 98°C for 15 s, primer annealing at 65°C for 30 s and extension at 72°C for 30 s, followed by a final extension 72°C for 5 min. Samples were chilled at 4°C for 5 min. After cool-down, 5 U of Exonuclease I (ExoI, NEB) were added, incubated at 37°C for 30 min followed by mixing 1.8x volume of Agencourt AMPure XP beads to the PCR/ExoI mix and purified according to manufacturer’s protocol. Samples were eluted in 20 µl ultra-pure water. After library preparation, samples were analyzed using the TapeStation 2200 DNA ScreenTape assay (Agilent) and the molarity of each library was determined by the average size of the peak maxima and the concentrations obtained from the Qubit fluorimeter (Thermo Fisher Scientific). Libraries were multiplexed by mixing the same molar concentration (2-5 nM) of each sample library, and library and sequenced using the Illumina HiSeq 2500 sequencing system using either 2×51 paired end reads for the 5S-rRNA control and *in vitro* experiments or 2×101 bp paired-end reads for all other samples. See Table S4 for read counts for all experiments presented in the manuscript.

## QUANTIFICATION AND STATISTICAL ANALYSIS

### Single clone expression level analysis

The average normalized fluorescence of mCerulean, and rate of production of mCherry, were calculated for each inducer concentration using the routine developed in (Keren et al., 2013), as follows:

mCerulean average normalized fluorescence: for each inducer concentration, mCerulean measurements were normalized by OD. Normalized measurements were then averaged over the N logarithmic-growth timepoints in the interval [T0, Tfinal], yielding:

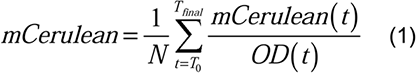

mCherry rate of production: for each inducer concentration, mCherry fluorescence at *T*_*0*_ was subtracted from mCherry fluorescence at *T*_*final*_, and the result was divided by the integral of OD during the logarithmic growth phase:

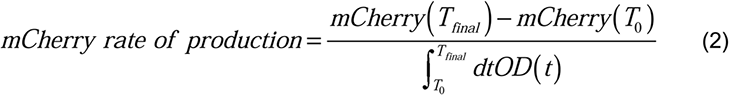

Finally, we plotted mCherry rate of production [(Zeevi et al., 2011)] as a function of averaged normalized mCerulean expression, creating dose response curves as a function of RBP-mCerulean fluorescence. Our choice for computing rate of production for mCherry stems from our belief that this observable best quantifies the regulatory effect, which is a function of the absolute number of inducer protein present (i.e RBP-mCerulean) at a any given moment in time. Data points with higher than two standard deviations calculated over mCerulean and mCherry fluorescence at all the inducer concentrations of the same strain) between the two duplicates were not taken into account and plots with 25% or higher of such points were discarded and the experiment repeated.

### Dose response fitting routine and K_d_ extraction

Final data analysis and fit were carried out on plots of rate of mCherry production as a function of averaged normalized mCerulean fluorescence at each inducer concentration. Such plots represent production of the reporter gene as a function of RBP presence in the cell. The fitting analysis and K_d_ extraction were based on the following two-state thermodynamic model:

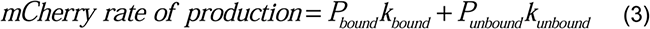

Here, the mCherry mRNA is either bound to the RBP or unbound, with probabilities *P*_*bound*_ and *P*_*unbound*_ and ribosomal translation rates *k*_*bound*_ and *k*_*unbound*_, respectively. The probabilities of the two states are given by:

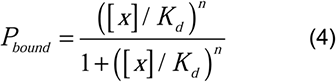

and

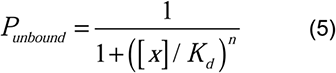

where *[x]* is RBP concentration, *K*_*d*_ is an effective dissociation constant, and n is a constant that quantifies RBP cooperativity; it represents the number of RBPs that need to bind the binding site simultaneously for the regulatory effect to take place. Substituting the probabilities into Eq. 3 gives:

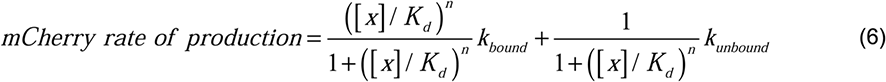

For the case in which we observe a down-regulatory effect, we have significantly less translation for high [*x*], which implies that *k*_*bound*_ ≪ *k*_*unbound*_ and that we may neglect the contribution of the bound state to translation. For the case in which we observe an up-regulatory affect for large [*x*], we have *k*_*bound*_ ≫ *k*_*unbound*_, and we neglect the contribution of the unbound state.

The final models used for fitting the two cases are summarized as follows:

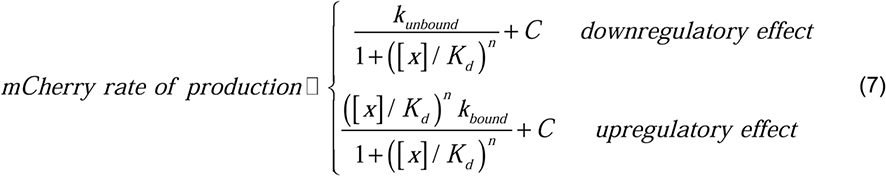

where *C* is the fluorescence baseline. Only fit results with *R*^2^ > 0.6 were taken into account. For those fits, *K*_*d*_ error was typically in the range of 0.5-20%, for a 0.67 confidence interval.

### SHAPE-Seq initial reactivity analysis

Illumina reads were first adapter-trimmed using cutadapt (Martin, 2011) and were aligned against a composite reference built from mCherry, E. coli 5S rRNA sequences, and PhiX genome (PhiX is used as a control sequence in Illumina sequencing). Alignment was performed using bowtie2 in local alignment mode (bowtie2 --local).

Reverse transcriptase (RT) drop-out positions were indicated by the end position of Illumina Read 2 (the second read on the same fragment). Drop-out positions were identified using an inhouse Perl script (can be provided upon request). Reads that were aligned only to the first 19 bp were eliminated from downstream analysis, as these correspond to the RT primer sequence. For each position upstream of the RT-primer, the number of drop-outs detected was summed.

To facilitate proper signal comparison, all libraries (16 total - including biological duplicates) were normalized to have the same total number of reads. For each library j and position x=1,…,L, we normalized the number of drop-outs Dj(x) according to:

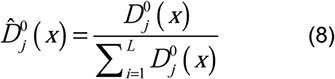

where *L* is the length of the sequence under investigation after RT primer removal. The reads as a function of position from the transcription start site (TSS) are supplied in Table S4.

### SHAPE-Seq Bootstrap analysis

To compute the mean read-ratio, reactivity, and associated error bars, we employed bootstrap statistics in a classic sense. Given *M* drop out reads per library, we first constructed a vector of length M, containing the index of the read # (1…M) and an associated position x per index. Next, we used a random number generator (MATLAB) and pick a number between 1 and *M, M* times to completely resample our read space. Each randomly selected index number was matched with a position *x.* The length *x* was obtained from the matching index in the original non-resampled library 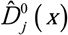. We repeated this procedure 100 times to generate 100 virtual libraries from the original 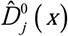 to generate 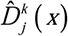, where k = {1..100}.

### SHAPE-Seq Signal-to-noise (read-ratio) computation

For each pair of NAI-modifed and umodified (DMSO) resampled libraries for a particular sample s 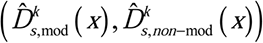, we computed the SHAPE-Seq read-ratio for each position *i* to generate a read-ratio matrix as follows:

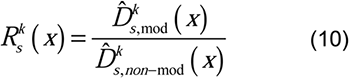

where the read-ratio is a signal-to-noise observable defined for each individual nucleotide. To obtain the mean read-ratio vector and associated standard errors, we computed the mean and standard deviation of the read-ratio per position as follows:

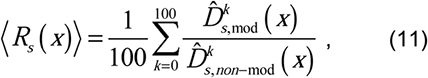

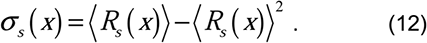

### SHAPE-Seq reactivity computation

The literature has several redundant definitions for reactivity, and no consensus on a precise formulation (Aviran et al., 2011; Lucks et al., 2011; Spitale et al., 2015) The simplest definition of reactivity is the modification signal that is obtained above the background noise. As a result, we define the reactivity as follows:

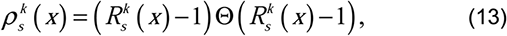

Where,

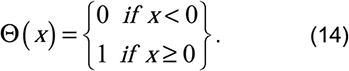

For the average reactivity score obtained for each position for a given sample s:

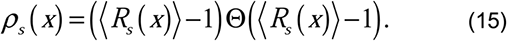

For the running-average reactivity plots shown in Figure 3, we used the following procedure. First, we computed an average reactivity per position based on two boot–strapped mean reactivity scores that were obtained from the two biological replicates. We then computed a running average 10 nt window for every position δ.

### SHAPE-Seq reactivity error bar computation

Error bars were computed in two steps. First, we computed the error-bar per nucleotide before running average as follows:

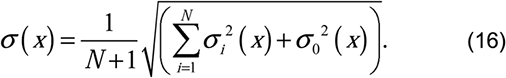

Where *σ*_*i*_ (*x*) corresponds to the boot-strapped sigma computed for position x of technical repeat *i*, while *σ*_0_ (*x*) is defined as the standard deviation at position *i* of the read ratio values for all *N* technical repeats. The error bar displayed for each position in the running average plot (Fig. 3A-B) were computed as follows:

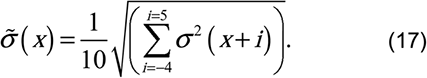

### SHAPE-Seq determining protected regions and differences between signals

To determine regions of the RNA molecules that are protected by the RBP, we employ a Z-factor analysis on the difference between the read-ratio scores. Z-factor analysis is a statistical test that allows comparison of the differences between means taking into account their associated errors. If Z > 0 then the two means are considered to be “different” in a statistically significant fashion (i.e. > 3*σ*). To do so, we use the following formulation:

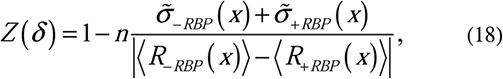

where *n* corresponds to the threshold of the number of *σ*’ *s* that we want to use to claim a statistically significant difference between two values of the mean. For our analysis we used *n = 3*. The regions that were determined to generate a statistically different mean reactivity values, and also resulted in a positive difference between the - RBP and +RBP cases (i.e. ⟨*R*_− *RBP*_ (*δ*) ⟩ − ⟨*R*_+ *RBP*_ (*δ*) ⟩) were considered to be protected and marked in a semi-transparent grey shading in Fig. 3 and Fig. 4.

### SHAPE-Seq structural visualization

For the structural visualization (as in Fig. 3C), the mRNA SHAPE-Seq fragment of PP7-wt_d=-29 construct was first folded *in silico* using RNAfold in default parameters. For visualization purposes, the RNAfold 2d structure prediction served as input for VARNA (Darty et al., 2009) and the SHAPE-Seq reactivity scores were used as colormap to overlay the reactivity on the predicted structure and to generate the structure image.

### Using the empirical SHAPE-Seq data as constraints for structural prediction

In order to predict more accurate structural schemes (Deigan et al., 2009; Ouyang et al., 2013; Washietl et al., 2012; Zarringhalam et al., 2012) we used the in vitro and in vivo SHAPE-Seq data as constraints to the computational structure prediction. This is done by taking the calculated reactivities of each sample, and computing a perturbation vector using RNApvmin of Vienna package (Lorenz et al., 2011) that minimizes the discrepancies between the predicted and empirically inferred pairing probabilities. Once the perturbation vector is obtained, we implement the Washietl algorithm (Washietl et al., 2012) in RNAfold to compute the inferred structure.

In order to calculate base-pairing probabilities for the structure determined by RNAfold with Washietl algorithm, the perturbation vector generated by RNApvmin is inserted as an additional input for RNAsubopt (-p 1000). A custom Perl script was used to calculate the resulted probability of pairing for each nucleotide based on the structural ensemble.

### SHAPE-Seq 5S-rRNA control

We first applied SHAPE-Seq to ribosomal 5S rRNA both in vivo and in vitro as a control that the protocol was producing reliable results (Kertesz et al., 2010; Spitale et al., 2015; Watters et al., 2016). We analysed the SHAPE-Seq read count by computing the “reactivity” of each base corresponding to the propensity of that base to be modified by NAI. Bases that are highly modified or “reactive” are more likely to be free from interactions (e.g. secondary, tertiary, RBP-based, etc.) and thus remain single stranded. We plot in Fig. S4 the reactivity analysis for 5S rRNA both in vitro and in vivo. The data shows that for the in vitro sample (red signal) distinct peaks of high reactivity can be detected at positions which align with single stranded segments of the known 5s rRNA (RFAM id: RF00001, PDB id: 4V69) (Szymanski et al., 2002; Villa et al., 2009; Watters et al., 2016).

By contrast, the in vivo reactivity data (blue line) is less modified on average and especially in the 9 central part of the molecule, which is consistent with these regions being protected by the larger ribosome structure in which the 5S rRNA is embedded (Dinman, 2005). The reactivity scores obtained here for both the in vitro and in vivo samples (Fig. S4B) are comparable to previously published 5S-rRNA reactivity analysis (Deigan et al., 2009; Szymanski et al., 2002; Watters et al., 2016).

### Tandem cooperativity fit and analysis

To estimate the degree of cooperativity in RBP binding to the tandem binding site, we used the following 4-state thermodynamic model:

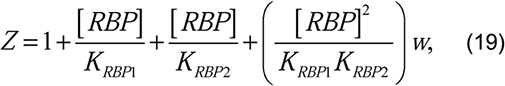

where *K*_*RBP1*_ and *K*_*RBP2*_ are the dissociation constants measured for the two single-binding-site variants, *[RBP]* is the concentration of the RNA binding proteins, and *w* is the cooperativity factor.

In a four state model, we assume four potential RNA occupancy states. No occupancy - receiving the relative weight 1. A state with single hairpin bound by an RBP receiving either the weight 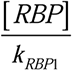 or the weight 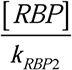 depending on whether the 5’ UTR or gene-header states are occupied respectively.

Finally, for the state where both hairpins are occupied we have the generic weight 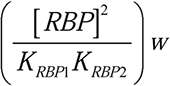, which takes into account also a potential interaction between the two occupied states, which can be cooperative if *w* > *1* or anti-cooperative if *w* < *1*. No interaction is the case where *w = 1*.

Next, we compute the relevant probabilities for translation for each weight. We know that when the ribosomal initiation region hairpin is occupied translation cannot proceed, however, some translation can result (albeit via a lower rate), when the 5’ UTR hairpin is occupied. This leads to the following rate equation for protein translation:

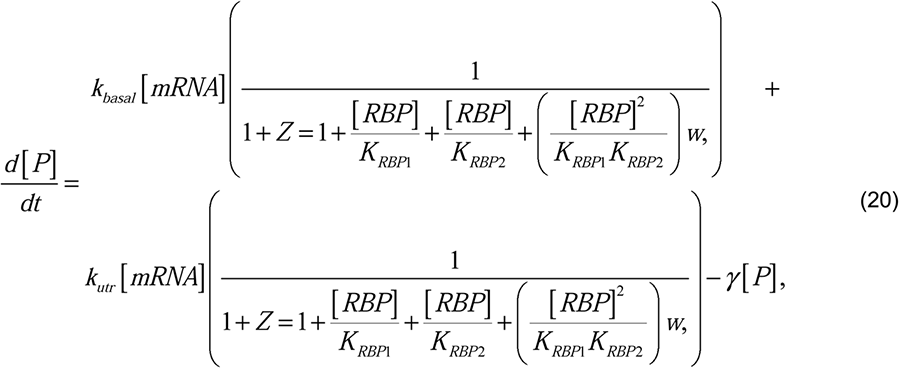

where *γ* is the protein degradation rate.

When measuring rate of production and given the stability of mCherry, the degradation rate of mCherry is negligible over the 1-2 hr range of integration that was used in 2. Since we normalized the basal levels of mCherry rate of production, 20 is reduced to the following fitting formula for the data:

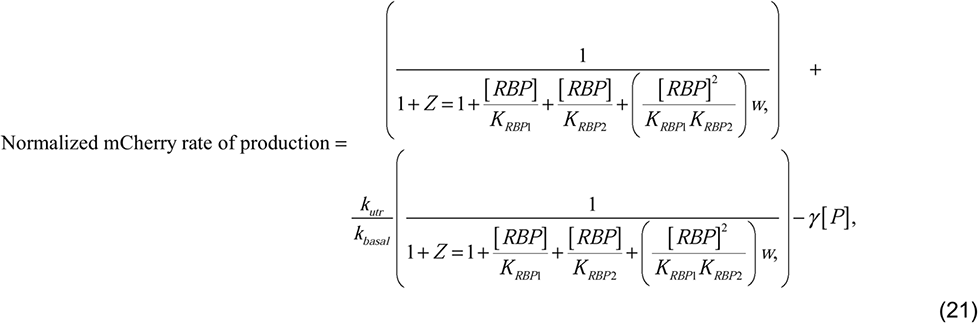

Finally, given our previous measurements for *K*_*RBP1*_ and *K*_*RBP2*_, this formula reduces to a two parameter fit for w and 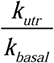. See Fig. S7 and Table S5 for the fits and associated fitting parameter details for 14 of 16 dose-response down-regulatory tandem data sets that were used in the analysis.

## DATA AND SOFTWARE AVAILABILITY

The SHAPE-Seq read data is available in doi:10.17632/wpknzwc2w2.1.

SUPPLEMENTARY TABLES TITLES:

1. Table S1: Reporter assay binding site cassettes
2. Table S2: Reporter assay RBP cassettes
3. Table S3: SHAPE-Seq sequences
4. Table S4: SHAPE-Seq reads
5. Table S5: Reporter assay tandem parameters

