## Supporting figures for "Synthetic 5’ UTRs can either up- or down-regulate expression upon RBP binding"

### Supplementary Information for:
### Synthetic 5' UTRs can either up- or down-regulate expression upon RBP binding

Noa Katz, Roni Cohen, Oz Solomon, Beate Kaufmann,
Orna Atar, Zohar Yakhini, Sarah Goldberg, and Roee Amit

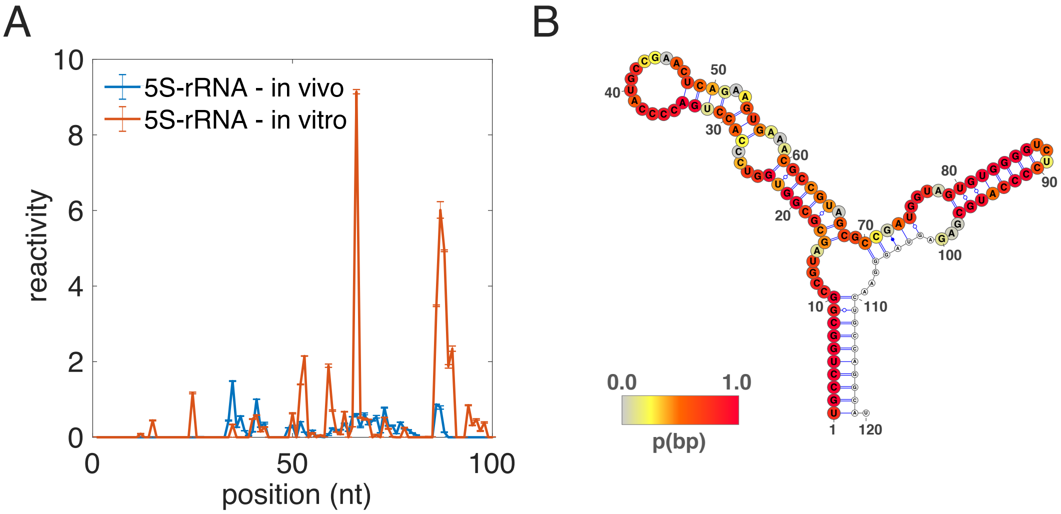


Figure S1: 5S-rRNA control. (A) reactivity scores for *in vivo* (blue) and *in vitro* (red) SHAPE-Seq measurements of 5S-rRNA. (B) 5S rRNA base-pairing probabilities were calculated using RNAfold and RNApvmin (by using the *in vitro* SHAPE-Seq data as constraints) and overlied as heatmap for each nucleotide on the known 5S rRNA structure (RFAM id: RF00001).

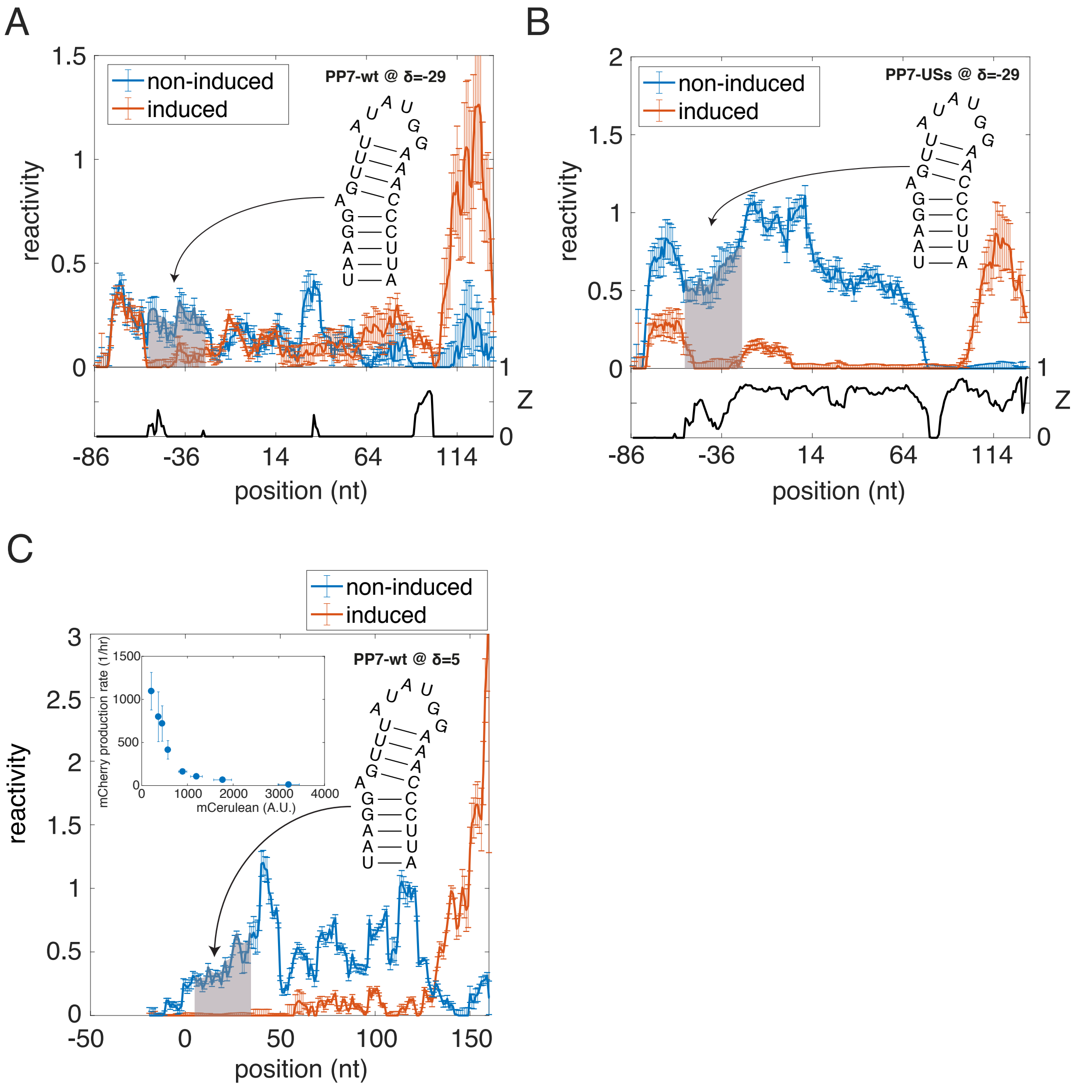


Figure S2: Induced vs non-induced plots for PP7-wt and PP7-USs *in vivo.* (A-B) Plots displaying the reactivities and Z-factor analysis (black) between the non-induced (blue) and induced (orange) strains for PP7-wt (A) and PP7-USs (B). Note the massive difference between the non-induced and induced states of PP7-USs in comparison to PP7-wt where only a small difference is observed in the vicinity of the binding site. (C) Plot comparing the non-induced (blue) to induced (orange) reactivity signals for PP7-wt when positioned at the ribosomal initiation region (δ=5). (Insets) Dose response plotted as mCherry production rate vs mCerulean fluorescence for PP7-wt (δ=5).

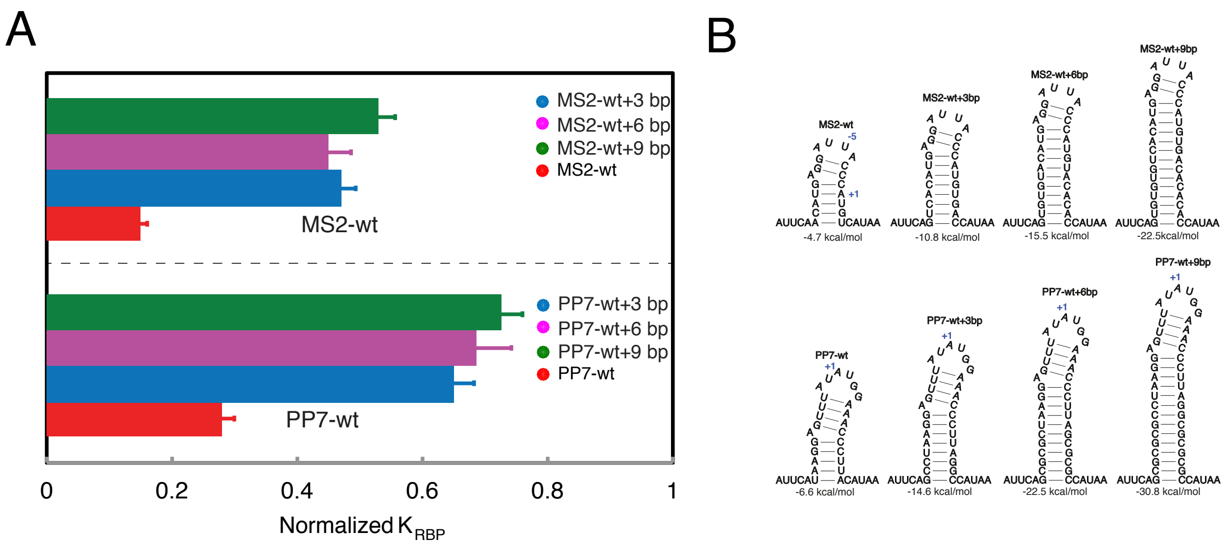


Figure S3: Normalized $K_{RBP}$s measured for long-stem variants. (A) $K_{RBP}$ calculated for all twelve constructs with extended stems for the corresponding RBPs (MCP or PCP). (B) Binding site schematic and free-energies of the PP7-wt and MS2-wt bindings that are augmented by longer stems.

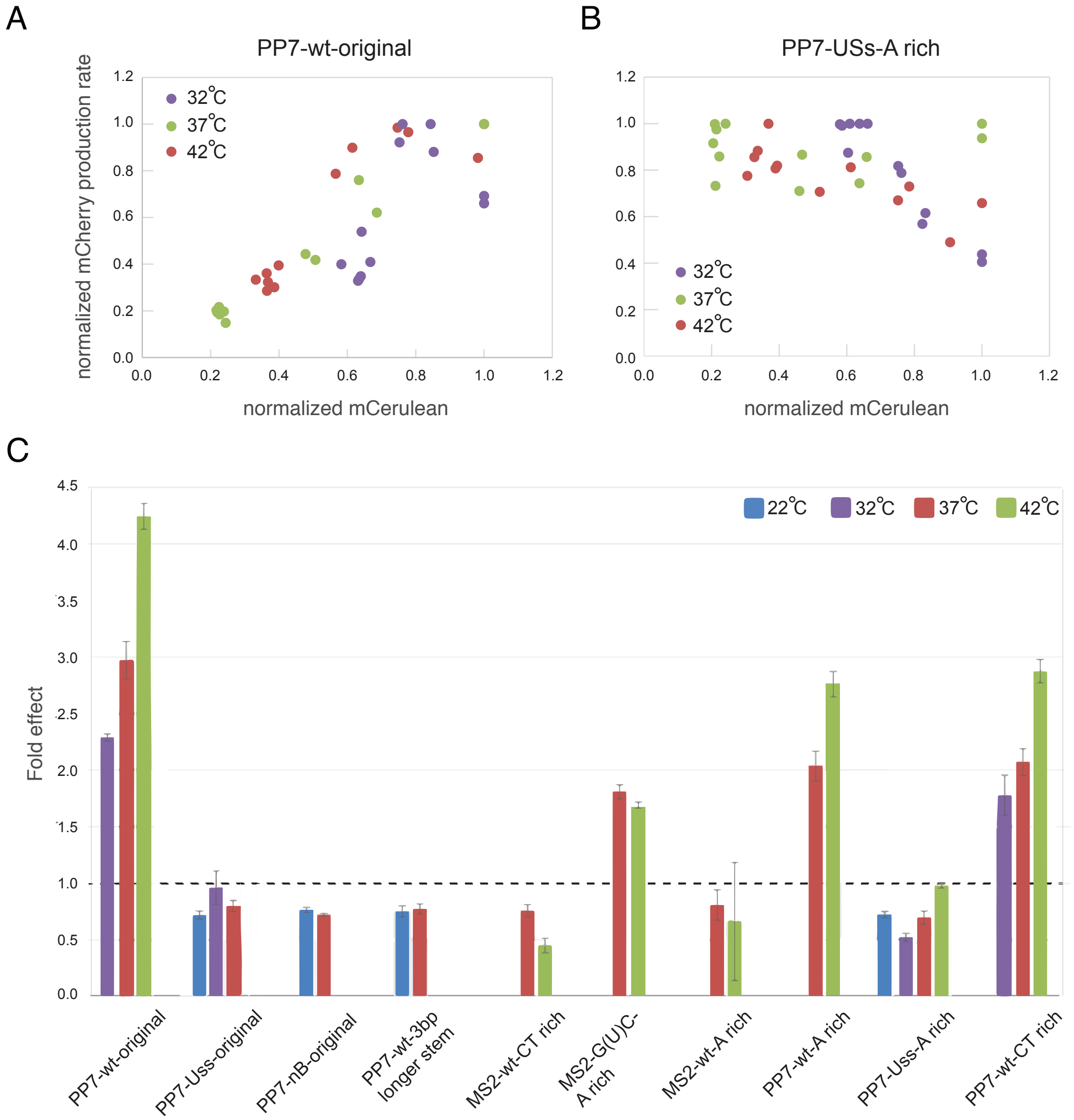


Figure S4: Temperature titration for selected variants. (A-B) Dose response functions measured at three ambient temperatures: 32, 37, and 42 °C for the following constructs: (A) PP7-wt (original), and (B) PP7-USs - A-rich. (C) Fold regulatory effects (up-regulation >1) for nine variants as a function of four temperatures exhibiting no change in the nature of the response (i.e., either down-regulating < 1, or up-regulating >1) as a function of temperature for all of the variants.

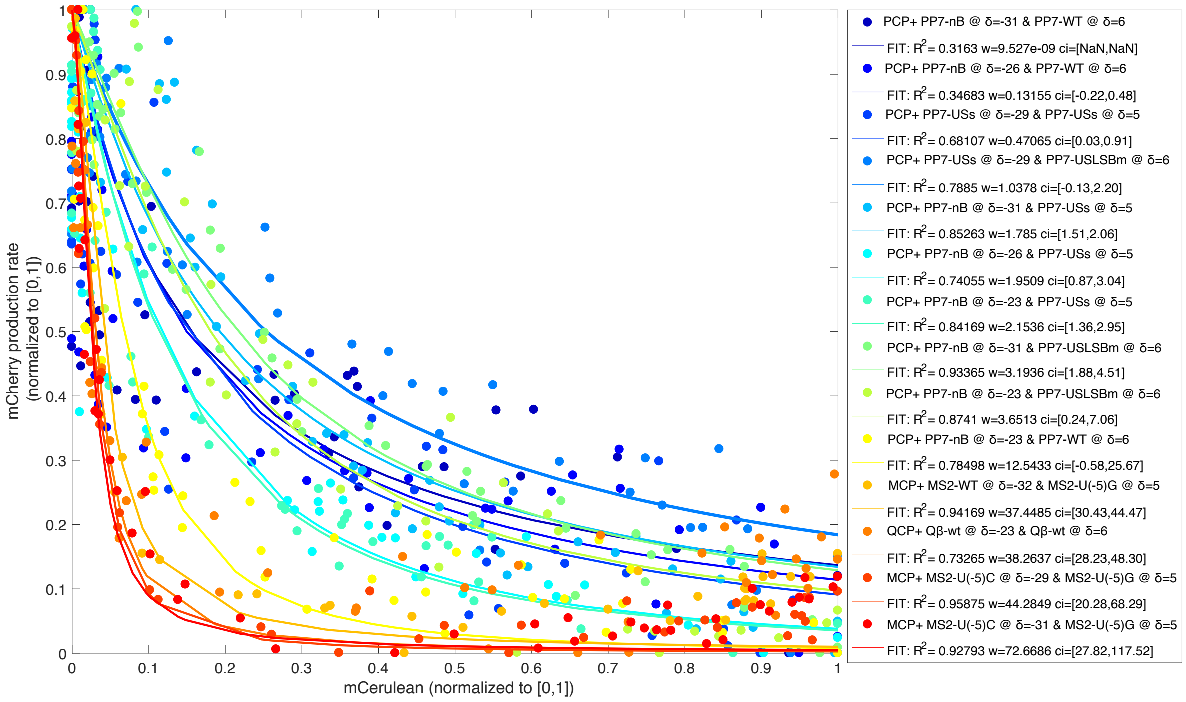


Figure S5: Tandem binding-sites dose response and associated fits. Dose responses and model fits for 14 of the 16 tandems whose cooperativity parameter was plotted in Fig. 7F.

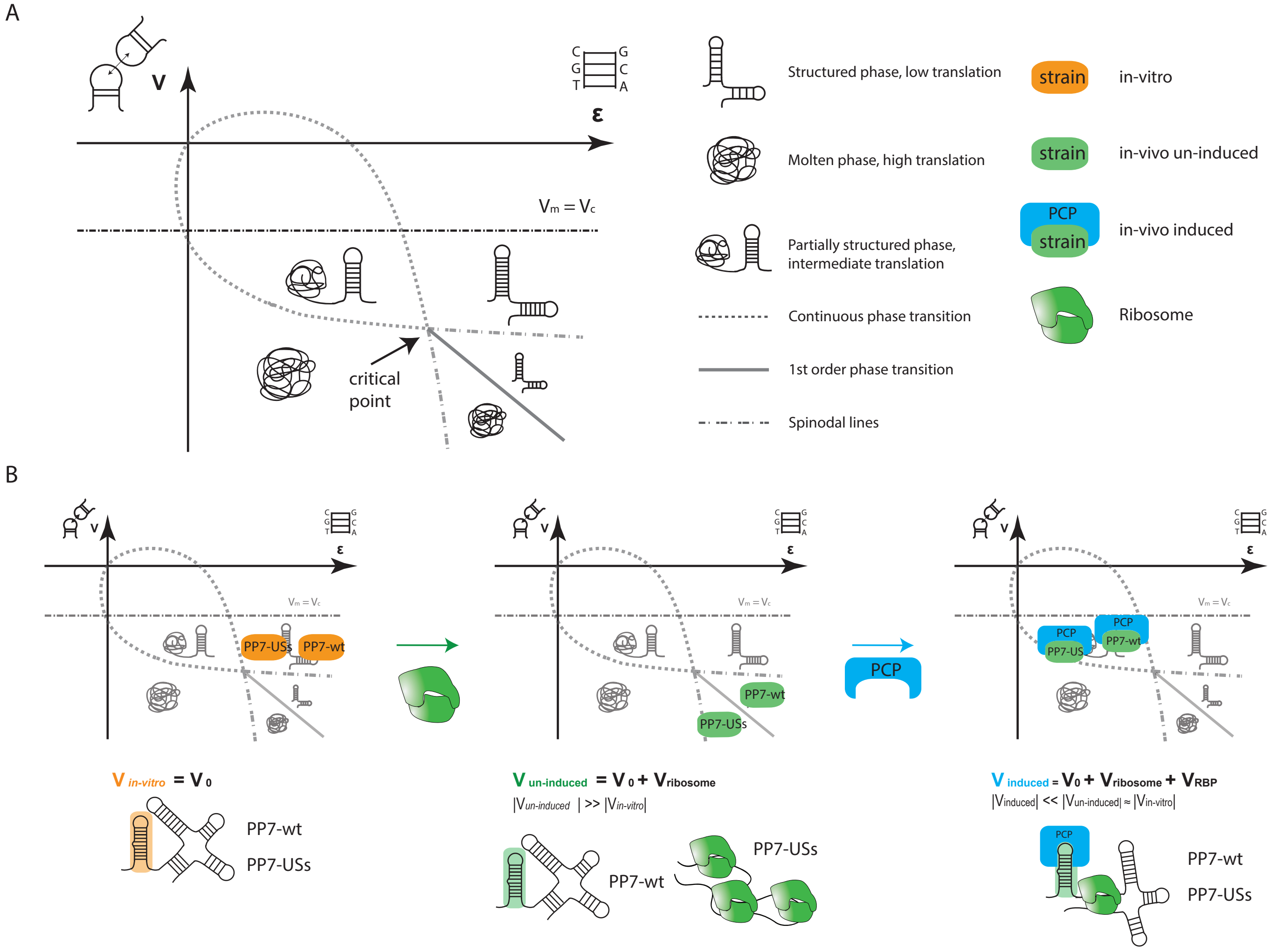


Figure S6: A schematic phase diagram providing an explanation for the two types of regulatory effects observed *in vivo*, despite having only one type of RBP-RNA interaction. (A) Reproduction of a mean-field phase diagram for RNA taking into consideration base-pairing energy ($\varepsilon$) and tertiary interactions between nucleotides that are unpaired ($\nu$) as originally published by (Schwab and Bruinsma, 2009). Negative $\nu$ values represent an attractive interaction. At values of $\nu$ below $\nu_o$ (dashed black line) and high $\varepsilon$ values, a structured phase is dominant. As $\varepsilon$ is reduced, a continuous phase transition occurs (dashed gray line) to a partially structured phase. Further reduction of $\varepsilon$ results in a second continuous phase transition from the partially-structured phase into the molten phase. In case $\nu$ is reduced below the critical point ($\nu_c$- gray arrow), a first order phase transition occurs (solid line) from the structured phase directly to the molten phase. (B) Utilizing the phase diagram in (A) to provide a model for the RBP-based regulation observed in our experiments. (*Left*) Hypothesized position of the PP7-wt and PP7-USs constructs (orange) on the phase diagram in the *in vitro* conditions. Here we assume a weak attractive tertiary interaction where $|\nu| < |\nu_c|$ and high $\varepsilon$ due to the constructs' sequences, which positions both constructs in the fully-structured phase. Note, that since PP7-wt has an additional base-pair with respect to PP7-USs, it is expected to have a slightly higher $\varepsilon$ value. (*Middle*) We assume that a subset of the types of interactions that contribute to $\nu$ also destabilize base-paired structures. This is based on the observation that in the analysis carried out by (Schwab and Bruinsma, 2009), a strong tertiary attractive interaction between un-base-paired elements inevitably destabilizes the base-paired structures. If so, the ribosome, which is known to destabilize secondary structures, may increase the probability of tertiary interaction between un-base-paired elements. In that case, the effect of the ribosome is to generate a stronger $|\nu|$ and lower $\varepsilon$, which may result in phase separation of the two constructs: PP7-wt remains structured on the right of the first order phase transition line, while PP7-USs is now in the molten state on the left of the first order phase transition line. (*Right*) Finally, we have previously shown using *in vitro* SHAPE-seq that PCP both stabilizes the binding-site structure, and simultaneously protects flanking regions from modifications (Katz et al., 2018). This implies that in the context of this model the RBP can both reduce $|\nu|$ below $|\nu_c|$, as well as alter $\varepsilon$ of the molecule. In this case, it is possible that the bound RBP repositions both PP7-wt and PP7-USs in the semi-structured phase.
